# Tracking live-cell single-molecule dynamics enables measurements of heterochromatin-associated protein-protein interactions

**DOI:** 10.1101/2023.03.08.531771

**Authors:** Ziyuan Chen, Melissa Seman, Ali Farhat, Yekaterina Fyodorova, Saikat Biswas, Alexander Levashkevich, P. Lydia Freddolino, Julie S. Biteen, Kaushik Ragunathan

## Abstract

Visualizing and measuring molecular-scale interactions in living cells represents a major challenge, but recent advances in microscopy are bringing us closer to achieving this goal. Single-molecule super-resolution microscopy enables high-resolution and sensitive imaging of the positions and movement of molecules in living cells. HP1 proteins are important regulators of gene expression because they selectively bind and recognize H3K9 methylated (H3K9me) histones to form heterochromatin-associated protein complexes that silence gene expression. Here, we extended live-cell single-molecule tracking studies in fission yeast to determine how HP1 proteins interact with their binding partners in the nucleus. We measured how genetic perturbations that affect H3K9me alter the diffusive properties of HP1 proteins and each of their binding partners based on which we inferred their most likely interaction sites. Our results indicate that H3K9me promotes specific complex formation between HP1 proteins and their interactors in a spatially restricted manner, while attenuating their ability to form off-chromatin complexes. As opposed to being an inert platform or scaffold to direct HP1 binding, our studies propose a novel function for H3K9me as an active participant in enhancing HP1-associated complex formation in living cells.

## INTRODUCTION

Genetically identical cells can exhibit different phenotypes due to covalent modifications of DNA packaging proteins called histones (1). These modifications give rise to heritable changes in gene expression, ultimately dividing the genome into two distinct states based on their intrinsic expression patterns: active euchromatin and silent heterochromatin (1). Heterochromatin comprises non-transcribed regions of the genome and is important for maintaining genome integrity, silencing repetitive DNA sequences, and preserving cell identity (2, 3). Certain histone modifications, like H3 lysine 9 methylation (H3K9me), are enriched within heterochromatin (4–6). The function of epigenetic modifications like H3K9me in establishing and maintaining gene silencing relies on the actions of specific histone modifier proteins (1). Histone modifiers often form large, dynamic multi-protein complexes with other accessory factors to regulate chromatin structure, genome organization, and transcription (2, 7). H3K9me acts as a binding platform for the recruitment of a conserved family of proteins called HP1, which play multiple roles in forming heterochromatin: HP1 proteins recruit histone modifiers that catalyze H3K9me deposition, enable the spreading of these modifications across large chromosomal regions, compact chromatin through oligomerization, and promote epigenetic inheritance following DNA replication (4, 8–12). Our current understanding is that H3K9me chromatin acts as a scaffold that transiently recruits HP1 and its binding partners to silence transcription (13, 14). However, HP1 proteins interact with diverse silencing and anti-silencing factors (15). The aberrant and simultaneous recruitment of these factors could lead to unproductive and competitive interactions that can jeopardize the integrity of gene silencing (16). How various histone modifier proteins interact with each other while sharing the same H3K9me substrate to assemble heterochromatin in living cells remains poorly understood.

Current methods for detecting protein-protein interactions are not sufficient for measuring real-time dynamical processes in living cells. Immunoprecipitation mass spectrometry detects protein-protein interactions, yet the results can depend on factors such as lysis conditions, salt concentrations, and protein abundance. As a result, these assays may not represent the full range of interactions that occur between proteins in living cells. Detecting protein-protein interactions after lysing cells also removes proteins from their native, crowded chromatin environment, so the *in vitro* properties of chromatin-associated factors can exhibit inconsistencies relative to how they behave in cells. For example, interactions with nucleic acids inhibit Swi6 nucleosome binding in vivo, whereas this binding is enhanced by nucleic acid *in vitro* (17). These disparities can be attributed to the significant variation in nucleic acid concentrations within the cellular environment: a large DNA excess can displace Swi6 from its binding site and promote protein turnover at sites of heterochromatin formation (17). Attempts to bridge the gap between *in vitro* and *in vivo* results with techniques such as Förster resonance energy transfer (FRET) and two-color imaging rely on protein-protein interactions that are infrequent and transient given the dynamic properties of chromatin-binding proteins. Additionally, FRET poses methodological challenges due to its limited working distance (< 10 nm) and the rare chances of spontaneous interactions between labeled molecules (18).

Live-cell super-resolution microscopy and single-molecule tracking are powerful tools for studying protein dynamics on the nanometer scale *in vivo* (19–21). Using photoactivatable fluorescent protein tags, we can track individual molecules to access the millisecond timescale interactions of histone modifiers with their chromatin substrates in living cells (17, 22, 23). The dynamics of these proteins in the cell nucleus represent the full range of possible interactions: protein diffusion is slowed by transient interactions and significantly reduced by binding to modified histones. Furthermore, proteins in cells bind not to individual nucleosomes but to large chromatin templates comprising hundreds of kilobases of modified nucleosomes decorated by a constellation of histone modifications. By leveraging recent advances in Bayesian inference methods, we analyze the complex trajectories that arise from interactions within a heterogeneous cellular environment (24, 25). This framework enables us to connect our biophysical measurements (a set of mobility states, each with an average diffusion coefficient) to protein-protein and protein-substrate interactions (a function of binding affinities) in living cells (17). Furthermore, by quantifying the probabilities of transitioning between each detected mobility state, we extend our measurements to derive on- and off-rates of protein-protein and protein-substrate interaction in cells (26).

Here, we use single-molecule tracking and Bayesian inference to measure chromatin-binding interactions in the model organism *Schizosaccharomyces pombe*, in which two HP1 orthologs, Swi6 and Chp2, bind to H3K9me (27–29). Despite their structural similarity and shared evolutionary origin, Swi6 and Chp2 are expressed at very different levels in the cell and play distinct roles in heterochromatin formation (16, 30). *In vitro*, Swi6 and Chp2 have similar tendencies to form dimers and oligomers (10, 16, 31, 32), but Swi6 binds to nucleosomes approximately 3-fold more strongly than Chp2 (31). Deletions of Swi6 and Chp2 have additive effects on epigenetic silencing, suggesting that Swi6 and Chp2 have distinct roles in establishing heterochromatin (16, 30, 31). However, it is unclear how their significant differences in expression levels and binding affinities affect how Swi6 and Chp2 interact with H3K9me in living cells.

Furthermore, the two HP1 proteins preferentially interact with different binding partners, yet it is still unclear how Swi6 and Chp2 specifically and selectively recruit their respective binding partners to sites of heterochromatin formation. Epe1 is a putative H3K9 demethylase (33) that interacts with Swi6 both *in vitro* and *in vivo* (34, 35) (**Figure 1**). On the other hand, the *S. pombe* SHREC complex, which consists of two major chromatin-modifying enzymes, Clr3 and Mit1, preferentially forms complexes with Chp2 (33, 36–38) (**Figure 1**). One explanation for these different associations is that Swi6 and Chp2 first form a complex with their respective partner proteins (Epe1 or Mit1 and Clr3) off chromatin, then search the genome, and ultimately bind at sites that are enriched for H3K9me. However, cells lacking Clr4 and H3K9me exhibit a significant loss in HP1-mediated protein interactions, suggesting that chromatin may play a causal role in enabling the assembly of heterochromatin-associated protein complexes in living cells (15). Hence, an alternative model is that HP1 proteins form complexes with their binding partners at sites of heterochromatin formation rather than off-chromatin.

**Figure 1.**
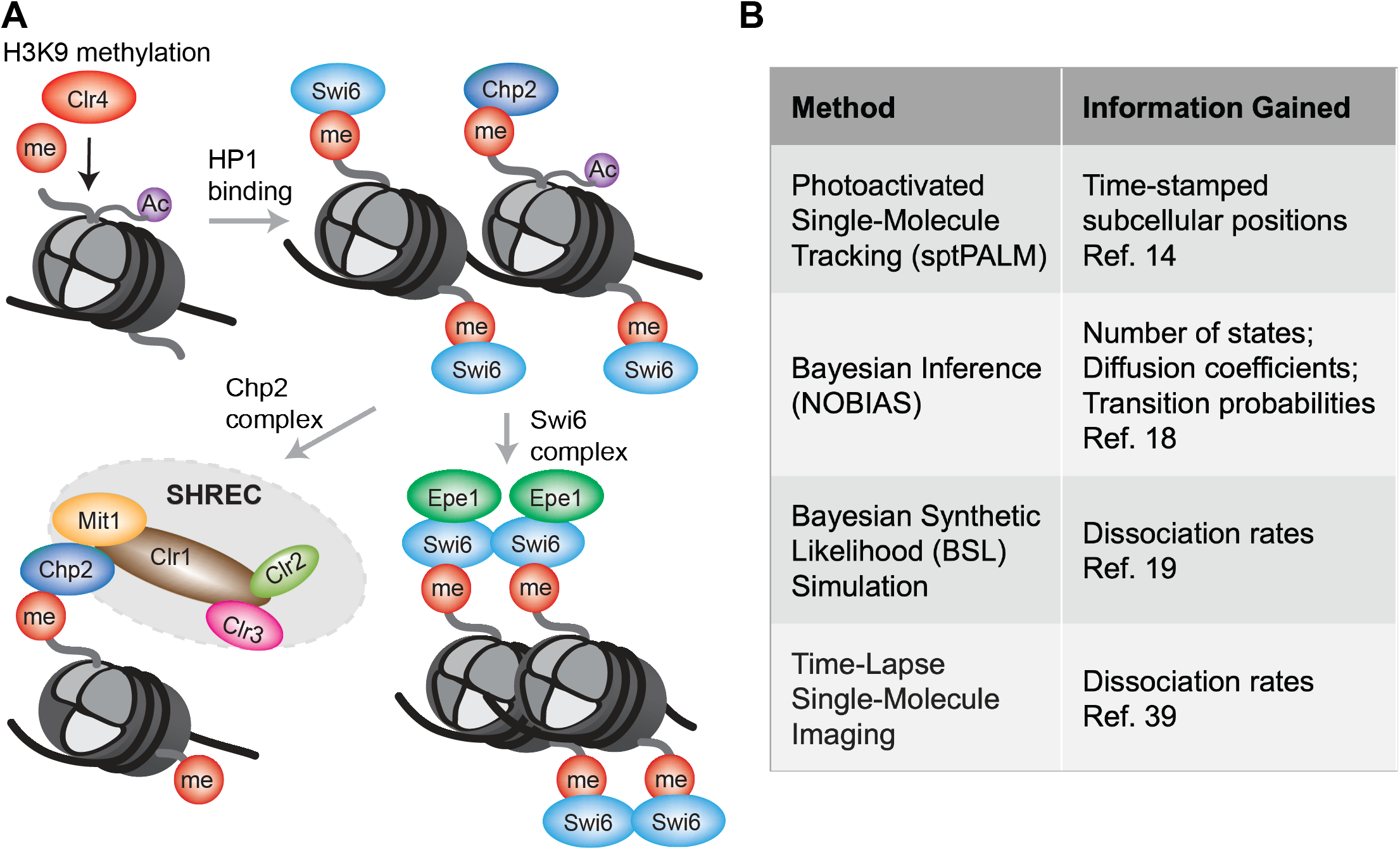
Single-molecule microscopy approaches for understanding interactions within HP1 protein complexes. **A**: current understanding of how HP1 protein complexes form on chromatin: Clr4 promotes H3K9 methylation at sites of heterochromatin; HP1 proteins Swi6 and Chp2 recognize H3K9me Swi6- and Chp2-dependent protein complex assembly at sites of heterochromatin. **B**: Methods used to access the interactions in HP1 protein complexes.

Here, we use live-cell single-molecule fluorescence microscopy to measure the dynamics and interactions of the HP1 proteins Swi6 and Chp2 and their primary interacting partners to infer how and where they form complexes within the context of native chromatin in living cells. Our results suggest that form complexes off-chromatin HP1 proteins form complexes with their binding partners at sites of H3K9me. As opposed to being an inert platform or scaffold to direct HP1 binding, our study proposes a potentially novel function for H3K9me in being an active participant that enhances the propensity of HP1 proteins to form complexes in living cells.

## RESULTS

### *S. pombe* HP1 orthologs, Swi6 and Chp2, exhibit distinct, non-overlapping biophysical states in living cells

We previously used single-molecule tracking to identify biophysical diffusive states that are associated with distinct biochemical properties of proteins in living cells (17). We measured the *in vivo* dynamics of Swi6, one of two HP1 proteins in fission yeast. We have determined that Swi6 has four distinct mobility states each of which maps to a specific biochemical property in cells (17). Here, we measured the mobility states of the second conserved HP1 protein, Chp2. We labeled the N-terminus of the endogenous copy of Chp2 with PAmCherry (PAmCherry-Chp2) but were unable to observe an appreciable number of photoactivation events for single particle tracking likely due to its low expression level. Instead, we inserted a second copy of an N-terminally labeled PAmCherry-Chp2 under the regulation of the thiamine-repressible promoters *nmt1, nmt41*, and *nmt81* (**Figure 2A, B**). We first ensured that *nmt81*-dependent expression of PAmCherry-Chp2 complements *chp2*Δ cells by measuring the silencing of a *ura4+* reporter inserted at the *mat* locus (*Kint2::ura4+*). If *ura4+* is silenced, cells grow on 5-fluoroorotic acid (EMMC+FOA) containing media and fail to grow on media lacking uracil (EMM-URA). We noted that *nmt81*-PAmCherry-Chp2 is functional and successfully restores *ura4+* silencing in *chp2*Δ cells (**Figure 2C**).

**Figure 2.**
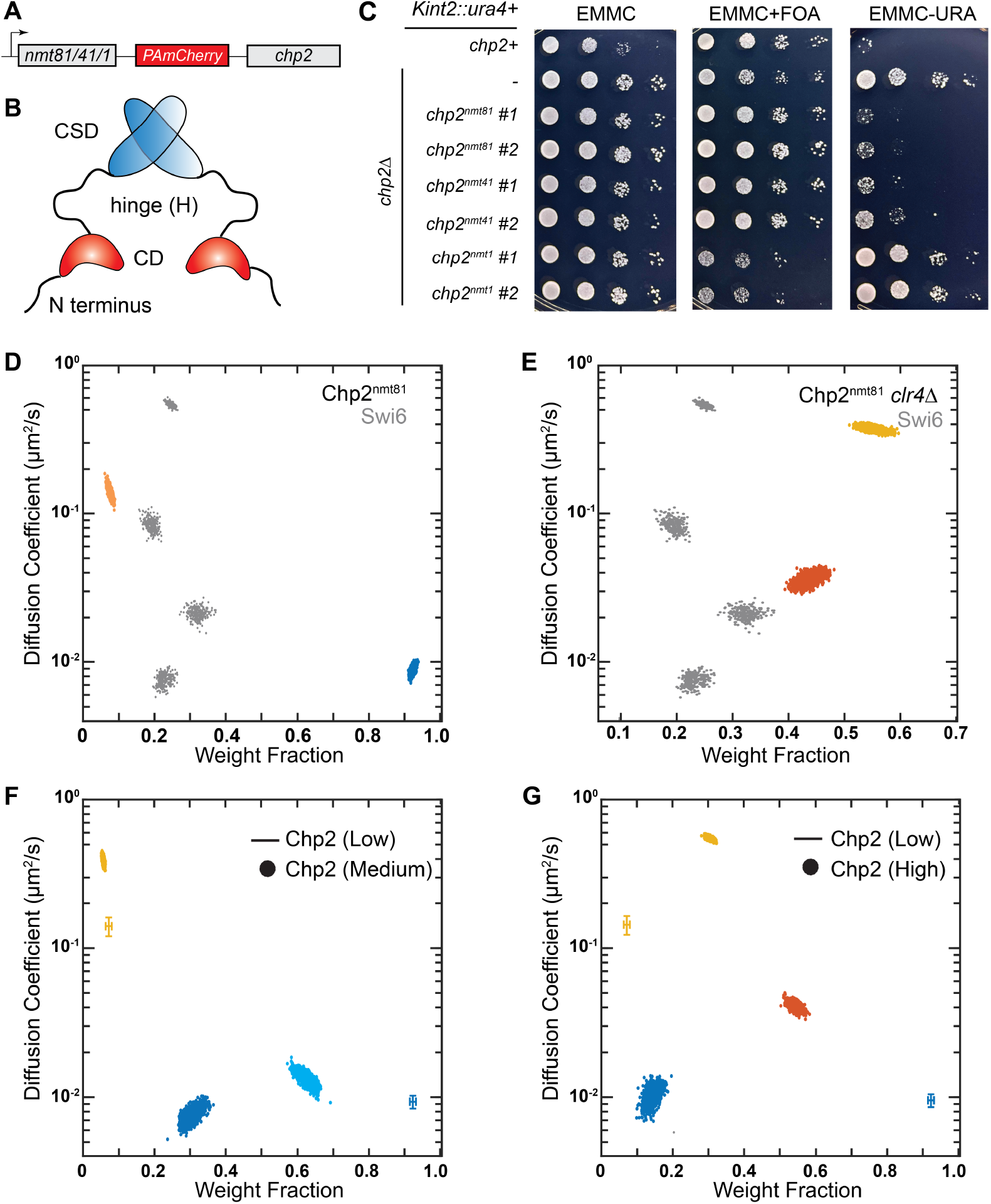
Single-molecule dynamics of PAmCherry-Chp2 at different concentrations. **A**: PAmCherry is fused to the N-terminus of Chp2 and expressed ectopically using a series of inducible promoters: *nmt1*, *nmt41*, and *nmt81*. **B**: Schematic of the Chp2 domains. CD: chromodomain (H3K9me recognition); H: hinge (nucleic acid binding); CSD: chromoshadow domain (dimerization interface). **C**: Silencing assay using an *ura4+* reporter inserted at the mat locus (*Kint2::ura4).* 10-fold serial dilutions of cells expressing Chp2 from different *nmt* promoters were plated on EMMC, EMMC +FOA, and EMM-URA plates. **D-E**: NOBIAS identifies two distinct mobility states for PAmCherry-Chp2^nmt81^. Each colored point is the average single-molecule diffusion coefficient of PAmCherry-Chp2 molecules in that state sampled from the posterior distribution of NOBIAS inference at a saved iteration after convergence in WT cells (**D**) and *clr4Δ* cells (**E**). Grey points are the previously reported PAmCherry-Swi6 single-molecule dynamics (24). **F-G**: NOBIAS identifies multiple mobility states for PAmCherry-Chp2^nmt41^ (medium expression, **F**) and PAmCherry-Chp2^nmt1^ (high expression, **G**). Each colored point is the average single-molecule diffusion coefficient sampled from the posterior distribution for PAmCherry-Chp2 at the indicated expression level. Colored crosses represent the data from PAmCherry-Chp2^nmt81^ (low expression; data in **D**).

Next, we tracked individual PAmCherry-Chp2 molecules expressed from the *nmt81* promoter in *S. pombe*. PAmCherry-Chp2 was briefly photoactivated with 405-nm laser light and imaged with 561-nm laser excitation light. We repeated this measurement until all PAmCherry-Chp2 molecules that can be activated were photobleached (see Methods). The activation-excitation-imaging cycle was repeated approximately 10 – 20 times for each cell, and the single molecules were localized and tracked in the recorded fluorescence movies with the SMALL-LABS algorithm (39). We model the motion of Chp2 molecules inside the *S. pombe* nucleus as a diffusive process and thus can assign diffusion coefficients to quantify the different mobility states associated with Chp2. We define a mobility state as a subpopulation of molecules with a distinct diffusion coefficient (D). In contrast to the four mobility states that we observed in the case of PAmCherry-Swi6 (a mixture of stationary and mobile molecules), nearly all PAmCherry-Chp2 proteins in *S. pombe* are stationary (**Figure S1B**). Hence, our live cell imaging data reveals a substantially different binding configuration between Swi6 and Chp2.

To investigate any potential heterogeneity in the dynamics within the observed static molecules, we applied NOBIAS, a nonparametric Bayesian framework that can objectively determine the number of mobility states giving rise to a single-molecule tracking dataset (25). We identified two mobility states associated with PAmCherry-Chp2: over 92% of the Chp2 molecules are in the low mobility state with an average diffusion coefficient, *D*_*α*_, *_Chp2_* = 0.007 µm^2^/s (**Figure 2D**) and around 7.5% of Chp2 molecules in a fast mobility state with *D*_*δ*_, *_Chp2_* = 0.13 µm^2^/s. NOBIAS analysis also provides the probability of a molecule transitioning between two mobility states within its trajectory: Chp2 molecules in the fast mobility δ state are much more likely to transition to the slower state compared with the reverse transition (**Figure 3A**). These weight fractions and transition probabilities indicate that Chp2 molecules predominantly occupy the slow *α* mobility state and only a very small proportion of Chp2 molecules occupy the fast *δ* state. This slow Chp2 motion is very different from the motion of the second *S. pombe* HP1 protein, Swi6: similar Bayesian analysis using the SMAUG package found that Swi6 molecules are distributed across four distinct mobility states (17).

**Figure 3.**
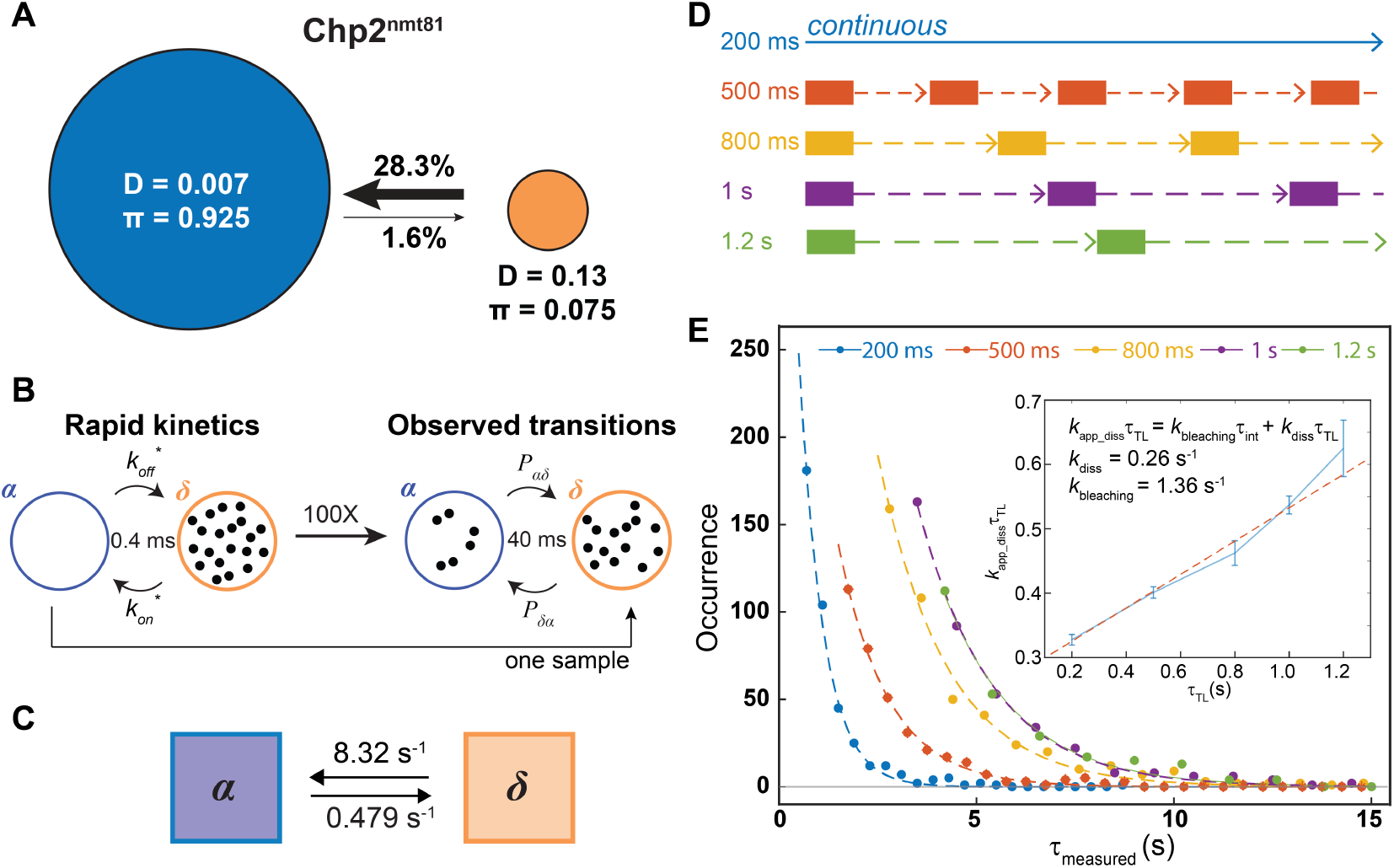
PAmCherry-Chp2 rate constants inferred from fine-grained chemical kinetic simulations and single-molecule time-lapse imaging. **A**: Inferred transition probabilities between the two mobility states of PAmCherry-Chp2^nmt81^ from single-molecule tracking (Figure 2D). Diffusion coefficients, *D*, in units of µm^2^/s and weight fractions, *π*, are indicated. Arrow widths are proportional to the transition probabilities. **B**: Fine-grained chemical kinetic simulations with Bayesian Synthetic Likelihood algorithm. The reaction on/off rate is proposed and simulated at a 0.4-ms time interval to calculate the likelihood based on transition probabilities from **A** at the 40-ms experimental imaging time interval. **C:** Inferred rate constants for PAmCherry-Chp2^nmt81^. **D:** Schematic of single-molecule time-lapse imaging. The time-lapse period, *r*_*TL*_, is the sum of the 200-ms integration time and the time delay. Five different time delays were introduced to access *r*_*TL*_ = 200, 500, 800, 1000, and 1200 ms. **E**: Dwell time distributions for PAmCherry-Chp2^nmt81^. The distributions are shown with fits to an exponential decay. Insert: linear fit (red dashed line) of *k*_*app*_*diss*_*r*_*TL*_vs. *r*_*TL*_, from which the dissociation rate constant, *k*_*diss*_, and the photobleaching rate constant, *k_bleaching_*, are obtained. Errors bars are the standard deviation of the exponential decay fitting.

To determine if the dominant slow mobility state of Chp2 corresponds to H3K9me-bound Chp2, we deleted Clr4, the only H3K9 methyltransferase in *S. pombe* (40, 41). In a *clr4*Δ background, the slowest PAmCherry-Chp2 mobility state is completely absent (**Figure 2E**). PAmCherry-Chp2 molecules in *clr4*Δ cells switch over to the fast mobility state consistent with Chp2 proteins moving around the nucleus in an unconstrained manner (weight fraction = 56%, *D_fast_* = 0.36 µm^2^/s). In addition, we observed a new mobility state that we did not previously detect in *clr4+* cells (weight fraction = 44%, *D_int_* = 0.03 µm^2^/s). The new mobility state most closely matches the chromatin sampling (*β* state) that we previously observed in the case of Swi6. Therefore, in the absence of H3K9me, Chp2 exhibits a substantial degree of binding to unmethylated chromatin. In contrast, only ∼10% of Swi6 molecules are in a chromatin sampling configuration with >60% of Swi6 molecules exhibiting fast, unconstrained diffusion in *clr4*Δ cells.

The overarching goal of our studies is to measure the biochemical properties of proteins and how they form complexes in the context of living cells. The appearance of a new mobility state in *clr4*Δ cells led us to hypothesize that Chp2 protein molecules that dissociate from H3K9me engage in a substantial degree of promiscuous off-target interactions. Unlike an *in vitro* experiment, we cannot change concentrations of proteins incrementally to determine binding affinities and specificities between proteins and their cognate ligands. Instead, we used two additional *nmt* promoter variants (*nmt41* and *nmt1*) to alter the overall Chp2 levels in wild-type cells. We used western blots to quantify the differences in expression across the three promoters. Chp2 expression driven by nmt41 relative to nmt81 promoters differs by around 50-fold and nmt1 relative to nmt81 promoters is approximately 1000-fold (**Figure S1A**). Hence, the promoter variants give us a substantial dynamic range in terms of Chp2 concentration to assess whether the mobility states we observed are in any way limited by substrate availability and how much excess protein is required to detect new mobility states (H3K9me nucleosomes in *clr4*+ cells).

The dynamics of the medium expressed (50-fold higher) PAmCherry-Chp2^nmt41^ are slightly increased compared to low expression *nmt81* driven PAmCherry-Chp2. The slow diffusive state in the low expressing PAmCherry-Chp2^nmt81^ cells splits into two states in the medium expression PAmCherry-Chp2*^nmt41^* cells with *D_slow1_* =0.005 µm^2^/s and *D_slow2_* =0.010 µm^2^/s (**Figure 2F**). In contrast, in the highly expressed PAmCherry-Chp2^nmt1^ (over 1000 fold compared to *nmt81*), we observe that only ∼15% of Chp2 is in the slow diffusive state; the remaining Chp2 molecules are in intermediate and fast states (**Figure 2G**). The new state we observed in the case of the high expression PAmCherry-Chp2 cells is comparable to Chp2 dynamics in a *clr4*Δ background*-* where Chp2 has no substrate. In this background, Chp2 molecules adopt a new mobility state that most closely resembles the chromatin sampling *β* state we observed previously in our Swi6 single particle measurements. Our *ura4+* reporter-based silencing assays revealed that PAmCherry-Chp2 expressed from an *nmt41* promoter (PAmCherry-Chp2^nmt41^) preserves *ura4+* reporter gene silencing whereas PAmCherry-Chp2 expressed from the high expression nmt1 promoter (PAmCherry-Chp2^nmt1^) disrupts silencing (**Figure 2C**) (16). This concentration-based effect may indicate that Chp2 outcompetes other chromodomain-containing proteins for a limiting amount of H3K9me substrate. At the extreme limit, the nmt1 promoter-driven expression of PAmCherry-Chp2 leads to the complete displacement of Swi6 from sites of heterochromatin formation. As expected, we observed labeled mNeongreen-Swi6 molecules uniformly distributed across the nucleus upon PAmCherry-Chp2^nmt1^ overexpression (**Figure S1D**). Hence, maintaining the equilibrium of Chp2 in a low diffusion state (H3K9me dependent) preserves its heterochromatin-associated silencing functions.

Swi6 and Chp2 both have a chromodomain that is responsible for H3K9me binding specificity (**Figure 2B**). We asked to what extent Swi6 competes with Chp2 to bind to H3K9me nucleosomes. We imaged PAmCherry-Chp2 in cells lacking the major HP1 protein, Swi6 (*swi6*Δ). Like in WT cells, the majority of Chp2 molecules in *swi6*Δ cells exhibit slow mobility. However, unlike WT cells, the slow population is split into two distinct slow mobility states with *D_slow1_* =0.005 µm^2^/s and *D_slow2_* =0.010 µm^2^/s, with only a very small portion in the fast state (**Figure S1C**). Similar to WT cells, the fast Chp2 state is very unstable: there is a high probability of transitioning from the fast state to one of the faster slow states (**Figure S2B**). The appearance of a split mobility state is likely because deleting Swi6 disrupts epigenetic silencing or because Swi6 makes unknown contributions to stabilizing Chp2 binding, although our results cannot distinguish between these two possibilities. Because nonparametric Bayesian approaches are known to have the potential for over-splitting (42), we validated the existence of the two slower states of Chp2^nmt81^ in *swi6*Δ cells and in PAmCherry-Chp2^nmt41^ cells by analyzing all of our Chp2 datasets using two other different single molecule tracking methods: Dirichlet process mixture models for single-particle tracking (DPSP) (43) and Spot-On (44). Both analysis methods capture a similar increase in dynamics and significant heterogeneity in the low mobility state tracks as NOBIAS (**Figure S1**).

### Chp2 dissociates faster from the H3K9me site *in vivo* than *in vitro*

The preponderance of the stationary H3K9me-binding state for PAmCherry-Chp2^nmt81^ implies that our high-resolution single molecule tracking measurements may overestimate Chp2 dissociation rates due to photobleaching. This parameter is crucial to determine Chp2 binding kinetics *in vivo* and determine the extent to which such measurements correlate with *in vitro* assays. We estimated the dissociation rate using two approaches: 1) single-molecule tracking followed by a Bayesian Synthetic Likelihood (BSL) simulation (26). This simulation-based approach has the benefit of not being affected by experimental time-resolution limits; and 2) single-molecule time-lapse imaging at different time intervals to ensure that photobleaching did not lead to an overestimation of the dissociation rate (45).

To infer the rate constants of transitions based on the NOBIAS transition matrices, we used a Bayesian Synthetic Likelihood algorithm, which has previously been applied to assess Swi6 dynamics (17, 26). At each step, we simulated the experimental outcome of the transitions with 0.4 ms time steps 2000 times for a set of rate constants (**Figure 3B**). BSL methods infer the most justifiable distribution of rate constants to estimate the value and uncertainty of the reaction rate. We applied the BSL method to analyze the output of the single-molecule tracking analysis and estimated that *k*_*diss*_ = 0.479 ± 0.005 s^-1^ (**Figure 3C**). We also experimentally determined Chp2 residence times and dissociation rates using single-molecule time-lapse imaging (Methods). Based on single-molecule time-lapse imaging at five different time intervals **(Figure 3D**), we calculated a Chp2-H3K9me disassociation rate of *k*_*diss*_ = 0.260 ± 0.018 s^-1^ and an average dwell time of 3.85 s (**Figure 3E**). In contrast, time-lapse imaging of Swi6 gives *k*_*diss*_ = 0.454 ± 0.051 s^-1^ and an average dwell time of 2.20 s (**Figure S2C**). Both experimental approaches (single-molecule tracking and single-molecule photobleaching) measured a dissociation rate more than 10-fold faster *in vivo* compared to previous *in vitro* measurements of Chp2 binding to H3K9me (9.6 ± 0.60 × 10^-3^ s^-1^ for me2 and 1.5 ± 0.27 10^-2^ s^-1^ for me3) suggesting that the complex in vivo chromatin environment substantially alters Chp2 chromatin binding properties (16). The *in vivo* time-lapse measurement of Chp2 and Swi6 disassociation rates reveal that Chp2 remains bound to H3K9me for a longer time than Swi6 *in vivo* suggesting that in fact, Chp2 binds to H3K9me chromatin with higher affinity. Our BSL analysis of single-molecule tracking data(17) also confirms that Chp2 dissociation rate (0.479 s^-1^) is ∼3-fold lower than that of Swi6 (1.27 s^-1^). Deleting Swi6 did not affect the residence time and dissociation rate of PAmCherry-Chp2^nmt81^ in *swi6*Δ cells which revealed a *k*_*diss*_ = 0.269 ± 0.031 s^-1^ and an average dwell time of 3.72 s (**Figure S2D**). The similarity in disassociation rates between PAmCherry-Chp2^nmt81^ in WT cells and *swi6*Δ cells indicates that although deleting Swi6 perturbs the overall dynamics of Chp2, it does not affect the intrinsic affinity between Chp2 and H3K9me chromatin.

### The anti-silencing factor Epe1 co-localizes with its HP1 binding partner primarily at sites of H3K9 methylation and exhibits limited off-chromatin dynamics

Having established the baseline dynamics of two major HP1 proteins in *S. pombe*-Swi6 and Chp2, we sought to determine how HP1 proteins interact with accessory factors to facilitate heterochromatin assembly. The putative H3K9me demethylase Epe1 is a major determinant of heterochromatin stability (33, 34, 46, 47). Epe1 directly binds to Swi6 and this interaction is essential for Epe1 recruitment to sites of H3K9me. Deleting Epe1 leads to both unregulated H3K9me spreading and increased epigenetic inheritance (33, 35). We labeled Epe1 at the C-terminus with PAmCherry (Epe1-PAmCherry-). To confirm if Epe1 molecules successfully localize at heterochromatin sites, we labeled Swi6 with mNeonGreen (mNeonGreen-Swi6) in cells and imaged the emission in the green channel (488-nm excitation) alongside Epe1-PAmCherry in the red channel (561-nm excitation). Overlaying mNeonGreen images with Epe1-PAmCherry super-resolution images indicates that Epe1 foci form at the periphery of Swi6-heterochromatin foci (**Figure 4A**).

**Figure 4.**
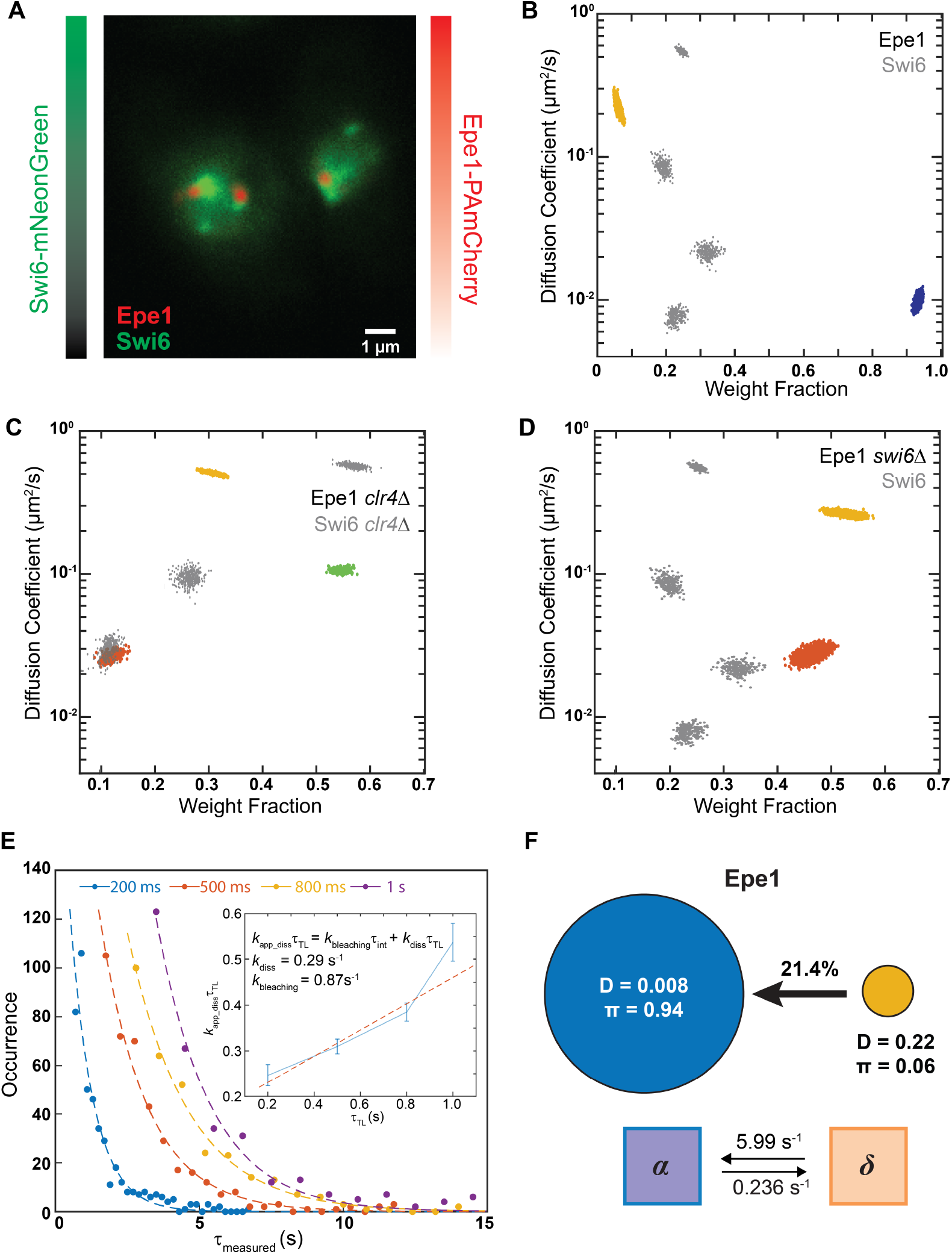
Single-molecule dynamics of Epe1 reveal that H3K9me reinforces the Swi6-Epe1 interaction on heterochromatin and suppresses off-site binding. **A**: Two-color imaging of cells expressing mNeongreen-Swi6 and Epe1-PAmCherry. Swi6 and Epe1 are expressed from their endogenous promoters. Green colorbar: Swi6-mNeonGreen intensities; Red colorbar: reconstructed Epe1-PAmCherry density map. Both color channels are normalized to the maximum pixel intensity. **B-D**: NOBIAS identifies distinct mobility states for Epe1-PAmCherry. Each colored point is the average single-molecule diffusion coefficient of PAmCherry-Chp2^nmt81^ molecules in that state sampled from the posterior distribution of NOBIAS inference at a saved iteration after convergence in WT cells (**B**), *clr4Δ* cells (**C**), and *swi6Δ* cells (**D**). Grey points are the previously reported PAmCherry-Swi6 single-molecule dynamics (24). **E**: Dwell time distributions for Epe1-PAmCherry expressed under its endogenous promoter. The distributions are shown with fits to an exponential decay. Insert: linear fit (red dashed line) of *k*_*app*_*diss*_*r*_*TL*_vs. *r*_*TL*_, from which the dissociation rate constant, *k*_*diss*_, and the photobleaching rate constant *k*_*bleaching*_ are obtained. Errors bars are from standard deviation of exponential decay fitting. **F:** Top: Transition probabilities between the two mobility states of Epe1-PAmCherry from **B**. Diffusion coefficients, *D*, in units of µm^2^/s and weight fractions, *π*, are indicated. Bottom: Inferred rate constants for Epe1-PAmCherry from the fine-grained chemical kinetic simulation.

To identify the mobility states associated with Epe1, we tracked single Epe1-PAmCherry molecules and inferred the number of mobility states, the diffusion coefficients, and the weight fraction for each Epe1 state. Since the interaction between Epe1 and Swi6 is direct, we expected to observe four mobility states similar to what we previously observed with Swi6. In contrast, we found that Epe1 has only two mobility states and that the predominant slower state (weight fraction, *π_slow_* ∼ 94%, *D_slow, Epe1_* = 0.008 µm^2^/s) (**Figure 4B**). Only ∼6% of Epe1 are assigned to a faster state with *D_fast, Epe1_* = 0.22 µm^2^/s. The transition probabilities indicate that transitioning from the fast state to the slow state is much more favored than the reverse transition (21% to 0.8%) (**Figure 4F**). These results suggest that in the presence of H3K9me, Epe1 preferentially remains in the H3K9me bound state presumably through its direct interaction with Swi6. To validate that the recruitment of Epe1 to sites of H3K9me (i.e., the slow state) is dependent on Swi6, we performed single-molecule tracking measurements of Epe1-PAmCherry in a *swi6*Δ background. As expected, we observed a complete loss of the slow state and the appearance of a new mobility state with a higher diffusion coefficient than the slowest state that we measured in WT cells. In addition, the weight fraction for the fast state, *π_fast_*, increases from 6% in the WT background to over 50% in *swi6*Δ (**Figure 4D**).

To determine the role that H3K9me might play in promoting complex formation, we performed Epe1-PAmCherry single molecule tracking measurements in *clr4*Δ cells. As expected, the previously observed Epe1 foci in wild-type cells disappear and Epe1-PAmCherry molecules in *clr4*Δ cells exhibit a diffuse distribution across the nucleus (**Figure S3C**). Also, we observed a complete loss of the slowest state given that neither Swi6 nor Epe1 can localize to sites of heterochromatin in the absence of their cognate H3K9me ligand (**Figure 4C**). Remarkably, we observed that Epe1 now exhibits three mobility states, and the diffusion coefficients of these states perfectly align with those of Swi6 (**Figure 4C**). The alignment in mobility states between Epe1 and Swi6 suggests that the two proteins can directly interact with each other and form off-chromatin complexes in the absence of H3K9me. The transition out of the fast state for Epe1 is 26 times higher than the transition into the fast state (**Figure 4F**) whereas this ratio decreases to 1.9 in *clr4*Δ cells (Figure S3B). These results support the idea that H3K9me chromatin shifts the complex formation equilibrium between Epe1 and Swi6 towards a chromatin-bound state.

Next, we performed time-lapse imaging to measure the Epe1 dissociation rate to compare how these rates differ relative to Swi6. We estimated that Epe1 dissociates from sites of heterochromatin formation at a rate that is *k*_*diss*_ = 0.288 ± 0.044 s^-1^ according to single-molecule time-lapse imaging with four time intervals (**Figure 4E**). BSL analysis of the single-molecule tracking transition matrix gave *k*_*diss*_ = 0.236 ± 0.003 s^-1^ (**Figure 4F**) or a dwell time of 4.24 s. These data suggest that Epe1 remains bound to heterochromatin for dwell times that are much longer than that of Swi6. These dissociation rate measurements suggest additional contributions beyond just the interaction between Epe1 and Swi6 which promote the persistent and stable association of Epe1 at sites of heterochromatin formation.

### Histone remodeler Mit1 and histone deacetylase Clr3 assemble into SHREC complex only at heterochromatin

Given our observation that Epe1 and Swi6 preferentially form complexes at sites of H3K9me and not off-chromatin, we wanted to determine the extent to which the principle of H3K9me-directed complex assembly might generalize to other HP1 protein complexes. The SHREC complex consists of a histone remodeler Mit1 and histone deacetylase (HDAC) Clr3 (**Figure 5A**)(36, 37). Unlike Epe1, which critically depends on Swi6 for its recruitment to heterochromatin, proteins that are part of the SHREC complex mostly form complexes with Chp2 (30, 37). The C-terminus of Chp2 forms a complex with the N-terminus of Mit1 and their interactions have been characterized using X-ray crystallography (37). This is further supported by studies of Swi6 purification followed by mass spectrometry in *chp2*Δ cells which reveals a precipitous loss of Mit1 from heterochromatin (15). In contrast, the recruitment of Clr3 depends both on HP1-dependent and HP1-independent interactions (30, 37, 48).

**Figure 5.**
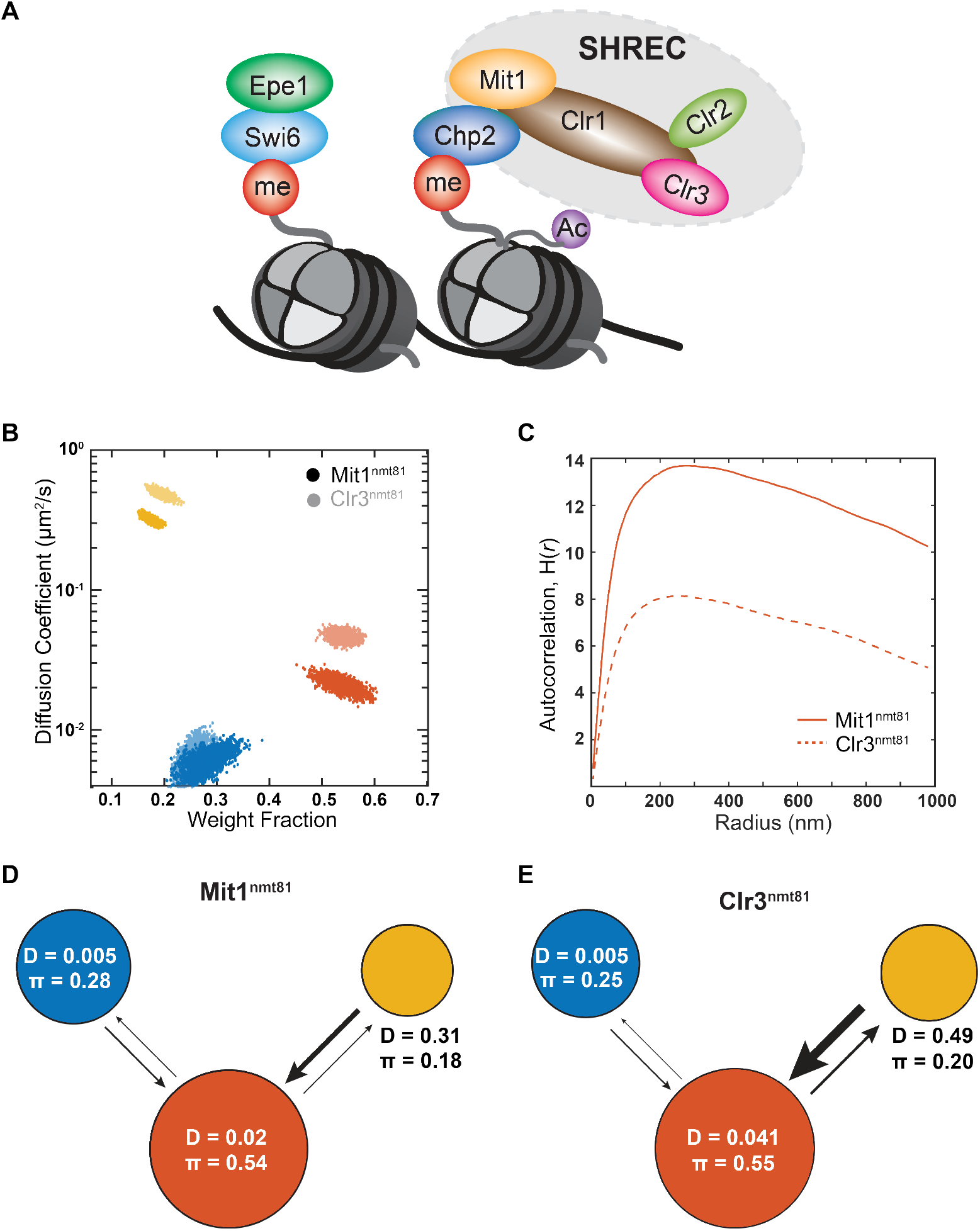
The two SHREC complex subunit proteins Mit1 and Clr3 only assemble at heterochromatin sites. **A**: Schematic of H3K9 methylated nucleosomes interacting with HP1 proteins and forming HP1 sub-complexes. Swi6 binds to Epe1; Chp2 interacts with the SHREC complex through Mit1. **B:** NOBIAS identifies three distinct mobility states for PAmCherry-Mit1^nmt81^ and PAmCherry-Clr3^nmt81^. Each point is the average single-molecule diffusion coefficient of PAmCherry-Mit1^nmt81^ molecules (solid colored points) or PAmCherry-Clr3^nmt81^ molecules (transparent colored points) in that state sampled from the posterior distribution of NOBIAS inference at a saved iteration after convergence. **C:** The intermediate state of Mit1^nmt81^(solid line) has a higher Ripley’s H(*r*) than the intermediate state of Clr3^nmt81^ (dashed line). Each autocorrelation plot is normalized with randomly simulated trajectories from the same state (Methods). **D-E** Transition probabilities between the three mobility states of PAmCherry-Mit1^nmt81^ (**D**) and PAmCherry-Clr3^nmt81^ (**E**) from NOBIAS. The arrow widths are proportional to the transition probabilities. Diffusion coefficients, *D*, in units of µm^2^/s and weight fractions, *π*, are indicated.

We previously determined that Chp2 exhibits two distinct mobility states and hence we extended our studies to identify the mobility states associated with its primary interacting partners - Mit1 and Clr3. We fused PAmCherry to the N-terminus of Mit1 and Clr3 and expressed the two fusion proteins using a thiamine-repressible *nmt81* promoter. We determined that PAmCherry-Mit1^nmt81^ preserved epigenetic silencing at the *mat* locus by using a *ura4+*-based silencing assay (**Figure S4A**). As previously described, the establishment of *ura4+* silencing leads to growth in FOA (EMMC+FOA) containing media and the lack of growth in media without uracil (EMM-URA). Our single-molecule tracking data for PAmCherry-Mit1^nmt81^ and PAmCherry-Clr3^nmt81^ reveals that both proteins exhibit three mobility states (**Figure 5B**). The diffusion coefficients for Mit1 and Clr3 only match each other for the slowest states (*D_slow_* = 0.005 µm^2^/s) with comparable weight fractions (28% for Mit1 and 25% for Clr3). Notably, the *D_slow_* values for these two proteins are again at levels similar to what we have observed in the case of other heterochromatin-associated factors (*D_slow_* of Swi6, Chp2, and Epe1). Reconstructed single-molecule fits density heatmap of Mit1 show that their high-density hotspots also exhibit spatial patterns that are similar to Chp2, Swi6, and Epe1 while Clr3 has a more widely dispersed pattern (**Figure S4C**).

We analyzed transition probabilities and calculated spatial autocorrelations for Mit1 and Clr3 based on our single-molecule tracking data. We noticed that Clr3 has a higher transition probability from the fast state to the intermediate state compared with Mit1 (**Figure 5D, E**). Spatial autocorrelation analysis is useful especially when combined with the state label NOBIAS provides for each step. We used the Ripley’s H function to determine the spatial overlap between Mit1 and Clr3 for different mobility states (49). A higher H(*r*) value indicates a higher clustering level at searching radius *r*, and Mit1 has a higher H function value than Clr3 in the intermediate state at all searching radii (**Figure 5C**). In contrast, there is little difference between the H functions for the slowest states of Mit1 and Clr3, indicating that the clustering levels of the slowest Mit1 and Clr3 molecules are similar (**Figure S4B**). In summary, the spatial auto-correlation analysis and single-molecule dynamic measurements for Mit1 and Clr3 suggest that the SHREC complex components preferentially co-localize only when H3K9me is present and both components are unlikely to form off-chromatin complexes.

Whether Chp2 and Mit1 recruit the HDAC module Clr3, to sites of H3K9me or the two SHREC complex components are recruited to the H3K9me site independently remains an open question (30, 37). We acquired and analyzed single-molecule tracking data for PAmCherry-Mit1 in *clr3Δ* cells, in which the number of diffusive states remains 3, and there is little change in the corresponding *D* and weight fraction for each state (**Figure S4D**), which means the binding of the remodeler module (Mit1) does not depend on the HDAC module (Clr3). In contrast, we found that the slow state of PAmCherry-Clr3 in *mit1Δ* cells has a decreased weight fraction and an increase in the diffusion coefficient associated with the slow state (*D_slow_* changes from 0.005 µm^2^/s to 0.010 µm^2^/s) (**Figure S4E**). These results suggest that Clr3 binding might depend on the successful binding and recruitment of Mit1. Alternatively, the deletion of Mit1 could directly affect heterochromatin stability.

### SHREC complex dynamics are affected by H3K9me

To test whether the slowest mobility state corresponding to Mit1 and Clr3 depends on H3K9me, we performed single-molecule tracking measurements in *clr4*Δ cells. In *clr4*Δ cells, Mit1 and Clr3 exhibit a substantial increase in the fastest mobility state (17.7% to 37.8% for Mit1 and 20.0% to 46.0% for Clr3) with a concomitant decrease in the slowest mobility state (28.4% to 10.7% for Mit1 25.5% to 13.0% for Clr3) (**Figure 6A,B**). However, unlike what we observed in the case of the two HP1 proteins—Swi6 and Chp2—or Epe1, the slowest mobility state is not fully eliminated for either PAmCherry-Mit1 or PAmCherry-Clr3 in *clr4*Δ cells. These results suggest that other mechanisms in addition to H3K9me are responsible for the slow mobility state of Mit1 and Clr3 (although more than half of its -bound state is determined by H3K9me). In Ripley’s H cluster analysis, for all steps of Mit1 and Clr3 in WT cells and *clr4Δ* cells, we notice a substantial decrease in the H(*r*) value for both proteins in the absence of Clr4, consistent with reduced clustering. (**Figure 6C**). Reconstructed localization maps of PAmCherry-Mit1 and PAmCherry-Clr3 in *clr4*Δ cells also show an overall unclustered spatial pattern for both proteins (**Figure 6D**).

**Figure 6.**
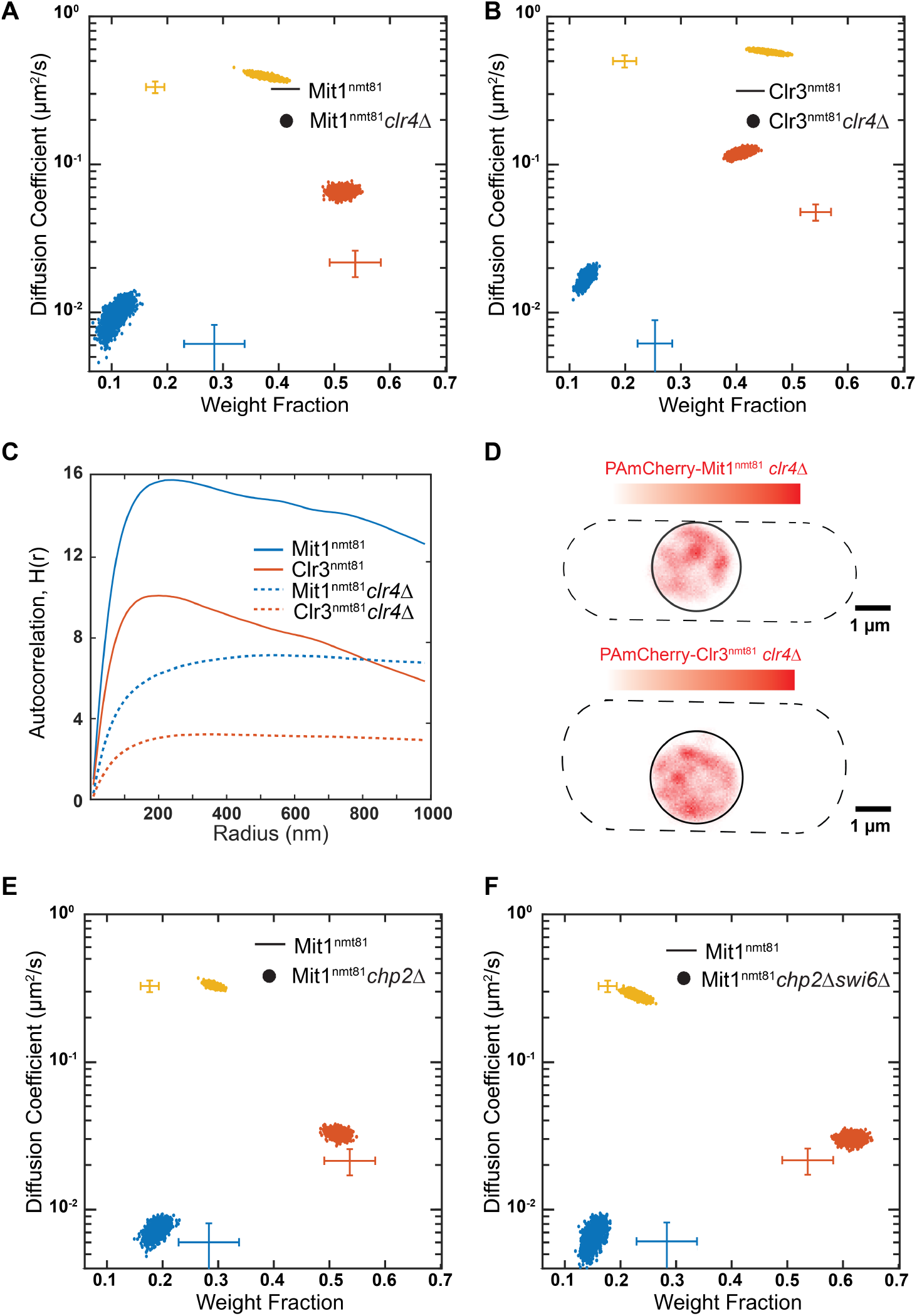
Chp2 associates with SHREC complex components only in the presence of H3K9 methylation. **A-B**: NOBIAS identifies three distinct mobility states for PAmCherry-Mit1^nmt81^ (**A**) and PAmCherry-Clr3^nmt81^ (**B**) in *clr4Δ* cells. Each colored point is the average single-molecule diffusion coefficient of molecules in that state sampled from the posterior distribution of NOBIAS inference at a saved iteration after convergence. The colored crosses show the data for PAmCherry-Mit1^nmt81^ and PAmCherry-Clr3^nmt81^ (Figure 5B). **C:** Ripley’s H analysis for steps from all states for Mit1^nmt81^ and Clr3^nmt81^ in WT cells and *clr4Δ* cells. Mit1^nmt81^ and Clr3^nmt81^ in *clr4Δ* cells have lower Ripley’s H(*r*) values than Mit1^nmt81^ and Clr3^nmt81^ in WT cells. **D**: Reconstructed single-molecule density map for PAmCherry-Mit1^nmt81^ (top) and PAmCherry-Clr3^nmt81^ (bottom) in *clr4Δ* cells. Dashed lines: approximate *S. pombe* cell outlines; solid circles: approximate nucleus borders. **E-F**: NOBIAS identifies three distinct mobility states for PAmCherry-Mit1^nmt81^ in *chp2Δ* cells (**E**) and *chp2Δ swi6Δ* cells (**F**). Each colored point is the average single-molecule diffusion coefficient of molecules in that state sampled from the posterior distribution of NOBIAS inference at a saved iteration after convergence. The colored crosses show the data for PAmCherry-Mit1^nmt81^ in WT cells (Figure 5B).

Next, we tested the extent to which the slow mobility state of Mit1 and Clr3 depends on the two HP1 proteins-Swi6 and Chp2. We acquired single-molecule tracking data of PAmCherry-Mit1 in *chp2Δ, swi6Δ,* and *swi6Δchp2Δ* cells. We analyzed the Mit1 single-molecule trajectories associated with each dataset and inferred the number of diffusive states and associated D and W values (**Figure 6E-F, S5A**). We notice that Mit1 in *chp2Δ* cells exhibits a substantial decrease in the bound state weight fraction compared to WT cells, but this decrease is less than what we observed in the case of *clr4Δ* cells (**Figure 6E**). In contrast, we observed a similar weight fraction for all three diffusive states of Mit1 in *swi6Δ* cells compared to Mit1 in WT cells (**Figure S5A**). These results suggest and indeed confirm that Chp2 is the primary HP1 protein interacting with Mit1. Finally, we observed that Mit1 dynamics in *swi6Δchp2*Δ produced an additive effect resulting in a further decreased bound state weight fraction compared with only *chp2Δ* cells (**Figure 6F**). In the absence of Chp2, Swi6 likely plays a compensatory role highlighting the potential for cross-talk between the two HP1 proteins.

For the HDAC Clr3, we acquired single-molecule tracking data for PAmCherry-Clr3 in *chp2Δ* and *swi6Δ* cells. In our analysis of Clr3 in *chp2Δ* cells (**Figure S5B**), we noticed a decrease in the bound state (similar to what we noted in the case of Mit1) in *chp2Δ* cells, which supports the hypothesis of Chp2-mediated SHREC recruitment. Deleting Swi6 (*swi6Δ* cells) (**Figure S5C**), also decreased the weight fraction for the Clr3 bound state, accompanied by an increase in the measured diffusion coefficient with *D_slow_* increasing from 0.005 µm^2^/s to 0.010 µm^2^/s which is well within the limit of our resolution. Our data shows that the stable nucleosome-bound state of the HDAC component Clr3, requires H3K9me, Chp2, and Mit1. We thus propose that the two modules of the SHREC complex (much like what we observed in the case of Epe1-Swi6) co-localize in the presence of H3K9me and with HP1 proteins only at sites of heterochromatin formation.

## DISCUSSION

In this work, we captured protein dynamics with super-resolution fluorescence imaging and analyzed single-molecule trajectories with Bayesian inference to investigate how heterochromatin-associated factors form complexes with their binding partners in living fission yeast cells (**Figure 1**). Our observations of the properties of heterochromatin-associated proteins in cells deviate in important and substantive ways from *in vitro* studies. Previous studies have shown that Swi6 binds to nucleosomes with a 3-fold higher affinity than Chp2 (31). In contrast, our data based on 1) the weight fractions of molecules in the H3K9me-dependent slow mobility state, 2) the transition rates of molecules between the free and bound states, and 3) time-lapse imaging to measure *k_off_* demonstrates that the majority of Chp2 molecules are H3K9me-bound and Chp2 binds with higher affinity to H3K9me chromatin. By altering Chp2 protein expression levels, we also reveal how Chp2 binds with exquisite specificity to H3K9me chromatin when expressed in limiting (and physiologically relevant) amounts. Hence, despite the two HP1 proteins having very similar domains, their different amino acid composition, especially within the nucleic acid-binding hinge domain, likely leads to different biochemical outcomes in cells. These results might explain why Chp2 is not easily displaced by Swi6 despite the endogenous expression levels of Chp2 protein being 100-fold lower than that of Swi6 in cells (16).

The binding properties of the two HP1 proteins—Swi6 and Chp2—serve as an important point of departure for our measurements on heterochromatin complex assembly in living cells. Epe1, a major anti-silencing factor that interacts with Swi6, exhibits only two mobility states, suggesting that Epe1 interacts with Swi6 exclusively at sites of H3K9me. These studies are consistent with our earlier observations that the addition of an H3K9me peptide dramatically increased the extent of binding between Epe1 and Swi6 *in vitro* (47). However, our previous *in vitro* biochemical analysis could not measure how Swi6 and Epe1 interact in cells in the context of H3K9me chromatin in living cells. Remarkably, our single-molecule measurements capture the exquisite specificity with which Swi6 and Epe1 form complexes in a heterochromatin-restricted manner, implying that the presence of H3K9me additionally suppresses off-chromatin interactions. Deleting Clr4 leads to Epe1 exhibiting three mobility states, and the Epe1 diffusion coefficients align with those of Swi6 in *clr4*Δ cells. Hence, Swi6 and Epe1 can exhibit pairwise interactions off-chromatin but the presence of H3K9me dramatically shifts the equilibrium populations toward a chromatin-bound state. The H3K9me chromatin-dependent enhancement also leads to Epe1 having longer dwell times at sites of heterochromatin relative to Swi6 (3.45 s versus 2.22 s).

We tested whether the principle of H3K9me enhancing complex formation could be extended to other proteins such as the chromatin remodeler, Mit1, and the histone deacetylase, Clr3, both of which form complexes with the second *S. pombe* HP1 protein, Chp2. Unlike Chp2, which has only two mobility states, Mit1 and Clr3 exhibit three mobility states. Both Mit1 and Clr3 exhibit mobility states with different diffusion coefficients and spatial autocorrelation functions, except for the slow state which we attribute to an H3K9me-bound fraction. These results suggest that Mit1 and Clr3, which are components of the SHREC complex, co-localize only at sites of H3K9me. Our results are consistent with recent structural work on SHREC complex proteins highlighting the special role that Chp2 plays in recruiting Mit1 to heterochromatin (37, 38).

Chromatin is largely thought to be a mere scaffold that recruits histone-binding proteins to particular locations in the genome (50). In contrast, our single-molecule measurements of heterochromatin proteins and their binding partners reveal a vital role for H3K9me as an enhancer of complex formation in living cells. Although the proteins whose properties we measured directly bind to each other and form pairwise interactions *in vitro,* we observe little off-heterochromatin co-localization when H3K9me is present. Our results reveal a dramatic shift in the equilibrium binding states induced by the presence of H3K9me, and these findings have important implications for the reconstitution and structural biology of heterochromatin-associated factors. Specifically, our measurements emphasize the need to explicitly include H3K9me chromatin substrates when describing models of how heterochromatin-associated factors form complexes both *in vitro* and in cells given its role in enhancing complex formation. Although the mechanisms of such enhancement are not well understood, it is likely that H3K9me recognition and binding trigger conformational changes that switch heterochromatin-associated proteins from a low-affinity to a high-affinity interaction state (47).

## MATERIALS AND METHODS

### Plasmids

All fluorescently tagged proteins were made with Gibson cloning. Strains with *nmt* promoters were constructed by modifying existing pDual plasmids (51).

### Strains

Most strains were constructed using PCR-based gene targeting approach (52). All strains with a PAmCherry fluorescent tag were made by constructing pDual vectors, containing the specified *nmt* promoter and the indicated protein(51). The plasmid was digested with the restriction enzyme, NotI, and the cut plasmid was transformed into *leu1-32* strains to select for growth on EMM-leu media (minimal media lacking leucine) as the pDual vector restores a functional *leu1*+ gene. Epe1-PAmCherry was made by using long oligos to tag Epe1. Deletions were made by PCR-based gene targeting approach or a cross followed by random spore analysis (53). All strains in this study are listed in Table S1.

### *S. pombe* live-cell imaging

Yeast strains were grown in standard complete YES media (US Biological, cat. Y2060) containing the full complement of yeast amino acids and incubated overnight at 32°C. For PAmCherry-Epe1 strains and Epe1 mutants under the control of the native Epe1 promoter, the seed culture was diluted into the same YES media and incubated at 25 °C with shaking to reach an OD_600_ ∼0.5. For strains with the nmt1, nmt41, or nmt81 promoter, the seed culture was diluted into EMMC media (FORMEDIUM, cat. PMD0402) containing the full complement of yeast amino acids and incubated at 30 °C with shaking to reach an OD_600_ ∼ 0.5. To maintain cells in an exponential phase and eliminate extranuclear vacuole formation, the culture was maintained at OD_600_ ∼0.5 for 2 days with dilutions performed at ∼12-hour time intervals (∼24-hour time interval for EMM media culture). Cells were pipetted onto a pad of 2% agarose prepared in EMM media and each agarose pad sample was imaged for less than 1 hour. *S. pombe* cells were imaged at room temperature with a 100× 1.40 NA oil immersion objective. The fluorescent background was decreased by exposure to 488-nm light (Coherent Sapphire, 377 W/cm^2^ for 20 – 40 s). A 406-nm laser (Coherent Cube 405-100; 1-5 W/cm^2^) was used for photoactivation (200 ms activation time) and a 561-nm laser (Coherent-Sapphire 561-50; 70.7 W/cm^2^) was used for excitation. Images were acquired at 40-ms exposure time per frame. The fluorescence emission was filtered to eliminate the 561-nm excitation source and imaged using a 512 × 512-pixel Photometrics Evolve EMCCD camera.

### Silencing assays

Strains containing the *ura4*+ reporter were grown overnight. Cells were equalized to 1OD/ml then four tenfold dilutions were spotted on minimal nonselective media (EMMC), minimal media lacking uracil (EMM-URA), or minimal media containing 5FOA (EMMC+FOA). 5-flouro-oratic acid (5-FOA) was added at a concentration of 1 g/L in EMMC+FOA plates. The plates were incubated at 32 °C for 3-4 days before imaging.

### Single-molecule trajectory analysis

Recorded PAmCherry single-molecule positions were localized and tracked with SMALL-LABS software(39). A mask of the nucleus of each cell was determined based on auto-florescence outside the nucleus in the 488nm bleaching step. Only the signal within the nucleus mask was analyzed. Single-molecule trajectory datasets were analyzed by a nonparametric Bayesian framework NOBIAS to infer the number of mobility states, the parameter for each state, and the transition between states (25). More than 1000 trajectories for each SPT dataset are put in the framework for robust analysis and to eliminate rare events. Reported parameters for each state are the posterior mean after the number of mobility states stabilizes, and reported uncertainty is the standard deviation from the posterior distribution. Some datasets were also analyzed with two publicly available SPT analysis software DPSP (43) and Spot-On (44). In DPSP analysis the chosen range of diffusion coefficients was 10^-3^ - 10 µm^2^/s. In Spot-On analysis, the number of components is set to 2 and 3 separately.

### Clustering Analysis for the Swi6 Distributions

The spatial pattern of each mobility state was investigated using the Ripley’s *K* function (49):

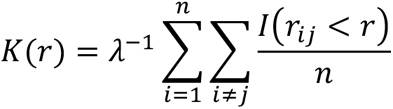

Where *r* is the search radius, *n* is the number of points in the set, *λ* is the point density and, *r*_i(_ is the distance between the *i*^th^ and *j*^th^ point. (*x*) is an indicator function. K(*r*) is further normalized to a Ripley’s H function:

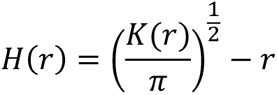

In this function, *H*(*r*) = 0 stands for a random distribution, *H*(*r*) > 0 means a clustered spatial pattern, and *H*(*r*) < 0 means a dispersed pattern. In the analysis, the nucleus was approximated as a circle to determine the area and perform edge correction (54). We calculated H(*r*) for each cell, and then we calculated an overall H(*r*) from the average of all cells weighted by the fits density.

To eliminate effects from the spatial correlation between single-molecule steps from the same trajectories, we simulated diffusion trajectories with similar confined area size, average track length, and overall density as experimental trajectories by drawing step lengths from the step size distribution of the corresponding experiment steps. This normalization is reported in previous work (17).

### Reconstructed single-molecule heatmap

For each cell, the nucleus and cell outlines were obtained from the fluorescence image of the nucleus and the phase-contrast image of the cell; these outlines were then approximated by a circle and a rectangle with circular caps, respectively. Every frame was analyzed by SMALL-LABS to identify single molecules, and the position and frame number of each single molecule were saved. To generate the reconstructed single-molecule heatmap for the cell, the pixel intensities after subtraction of the fitted offset in the appropriate diffraction-limited region about each single molecule were summed and the sum of all well-fit molecules was globally normalized.

### Fine-grained chemical rate constant inference

To infer the rate constants of transition for each of these proteins, we used a Bayesian Synthetic Likelihood algorithm similar to the one previously reported for the Swi6 rate constants (17). At each step, we began with a set of rate constants and their posterior density. We then proposed a potential new set in one of three ways based on a t-distributed random variable, *T*, with 10 degrees of freedom and the number of protein states *N*: we multiplied a random rate by exp(0.02*T*), we multiplied every rate by exp(0.02*T*⁄*N*^2^), or we multiplied a random pair of opposing rates by exp(0.01*T*). We simulated the experimental outcome of the transitions 2000 times for a set of rate constants. We used these simulations to calculate a likelihood distribution for the rate constants, multiplied the result by an improper Jeffries prior (*P*(*rate*) ∝ 1/*rate*), and compared the posterior density to that of the older set of rate constants using the standard Metropolis Hastings algorithm to decide whether to keep the old set of rates or accept the new one. We first simulated our system by dividing the experimental time (0.04 sec) into 100-time steps *δt* = 4×10^-4^ sec. We assumed that the sampled proteins were partitioned across states according to their equilibrium proportions based on their reported transition matrices from the NOBIAS single-molecule tracking analysis. We sampled from a binomial distribution with transition probability as given above and the number of molecules in a state to determine the number of molecules that transitioned. The posterior mean reaction rates were considered the final posterior rates. We report them with a 95% highest density credible interval.

### Single-molecule time-lapse imaging

We model the binding of Chp2 and H3K9me or Swi6 to Epe1 as a direct two-component association/disassociation reaction:

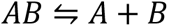

The measured residence time of each PAmCherry-Chp2 or PAmCherry-Epe1 molecule is estimated from the lifetime of the stationary fluorescence signal. *k_app_diss_* is acquired by fitting the probability distribution function, *P*, of the measured residence times, *τ_measured_*, to a single exponential decay function:

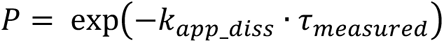

The measured apparent disassociation rate, *k_app_diss_*, consists of the true disassociation rate, *k_diss_*, and the photobleaching rate of the PAmCherry label, *k_bleaching_*; we separated these contributions by collecting data at multiple delay times to measure the photobleaching rate. For static molecules, we introduced a dark period with each time interval that we kept the integration time, *τ_int_*, the same and introduced different lengths of dark delay times, *τ_delay_*. In this way, the contribution of photobleaching was kept the same for different total time intervals, *r*_*TL*_ = *r*_*int*_ + *r*_*delay*_. We measured the residence time *r*_*measured*_ = (*n* − 1)τ_*TL*_ by counting the total number of sequential frames, *n*, in which the molecule was detected. Finally, the true disassociation rate, *k_diss_*, was estimated from a linear regression of the two-term relationship (45):

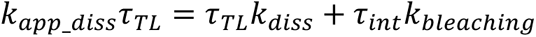

This linear regression also took the uncertainty of each data point from the exponential fitting into consideration and gave the final fitted slope *k*_*diss*_ and its uncertainty.

## Supporting information

Supplementary Figures and strains used in this study

## Acknowledgments

We thank Danesh Moazed for sharing the fission yeast strains used in this study. This work was funded by a National Science Foundation Understanding Rules of Life Award (1921677) to JSB, PF, and KR, and NIH award R35GM137832 to KR. This work used the Extreme Science and Engineering Discovery Environment (XSEDE) comet resource at the San Diego Supercomputing Center through allocation TG-MCB140220 to PF(55).

