## Supplementary Figures and strains used in this study for "Tracking live-cell single-molecule dynamics enables measurements of heterochromatin-associated protein-protein interactions"

### SUPPLEMENTARY FIGURE CAPTIONS

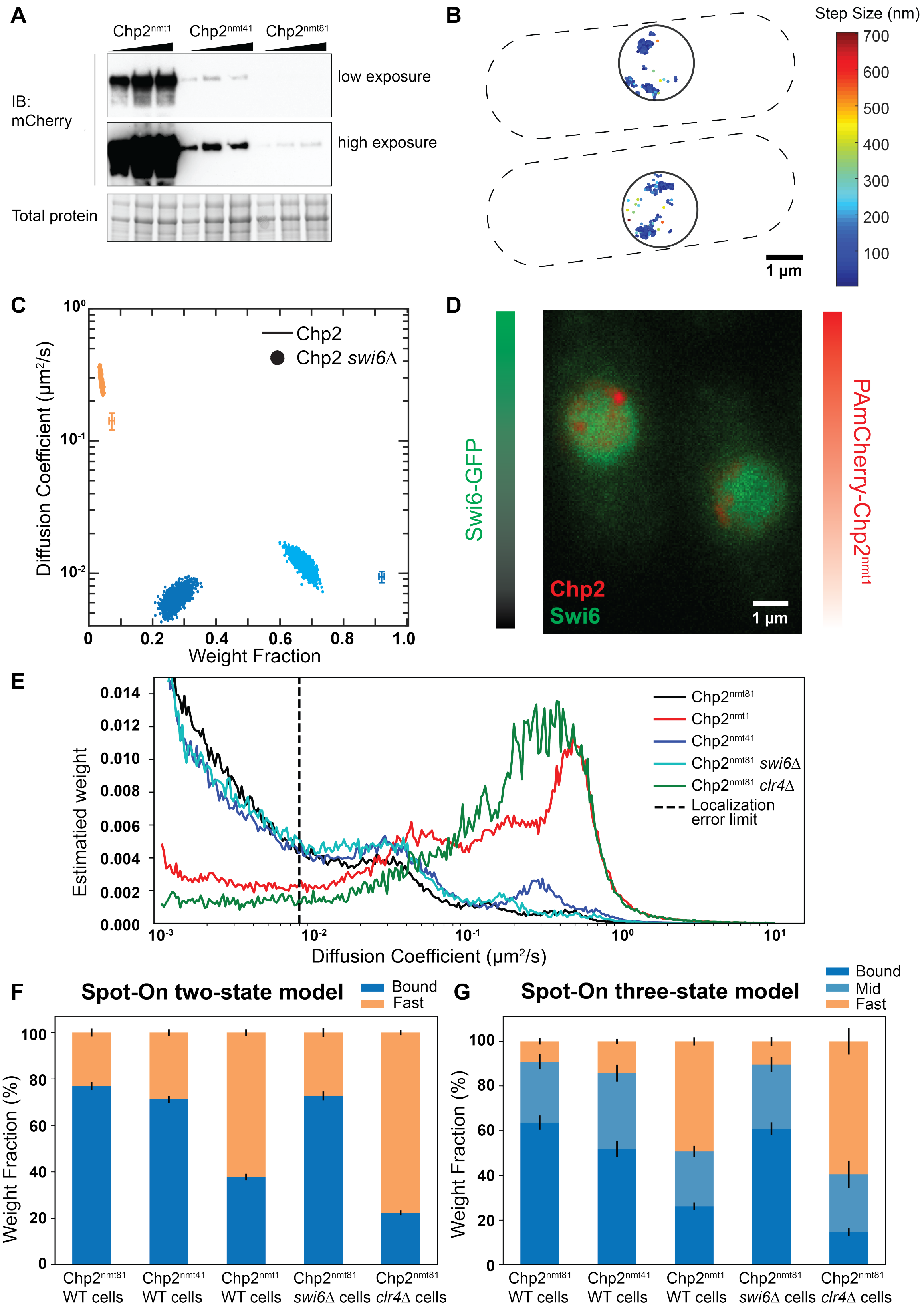


**Supplementary Figure 1. PAmCherry-Chp2 dynamics under different concentrations and environments with different analysis methods (caption on next page).**

**Supplementary Figure 1. PAmCherry-Chp2 dynamics under different concentrations and environments with different analysis methods. A**: The expression level of PAmCherry-Chp2$^{nmt1/nmt41/nmt81}$ is quantified by western blot against an mCherry antibody. The cross-reactivity of the mCherry antibody can specifically detect PAmCherry protein fusions. **B**: Single-molecule step size map for PAmCherry-Chp2$^{nmt81}$. Dashed lines: approximate *S. pombe* cell outlines; solid circles: approximate nucleus borders. **C**: NOBIAS identifies distinct mobility states for PAmCherry-Chp2$^{nmt81}$ in *swi6Δ* cells. Each colored point is the average single-molecule diffusion coefficient of molecules in that state sampled from the posterior distribution of NOBIAS inference at a saved iteration after convergence. The colored crosses show the data for PAmCherry-Chp2$^{nmt81}$ in WT cells (Figure **2D**). **D**: Two-color imaging of cells with Swi6-GFP expressed from the endogenous promoter and PAmCherry-Chp2$^{nmt1}$. Green colorbar: Swi6-GFP intensities; red colorbar: reconstructed PAmCherry-Chp2 density map. Both color channels are normalized to the maximum pixel intensity. **E**: Posterior distribution of diffusion coefficients of single-molecule trajectory datasets inferred from DPSP analysis (37). Vertical dashed line: Lower bound of detectable diffusion coefficients given the experimental localization error. **F-G**: Weight fractions of each mobility state for PAmCherry-Chp2$^{nmt81}$ single-molecule trajectories inferred from Spot-On analysis (38) with a two-state model (**F**) and a three-state model (**G**).

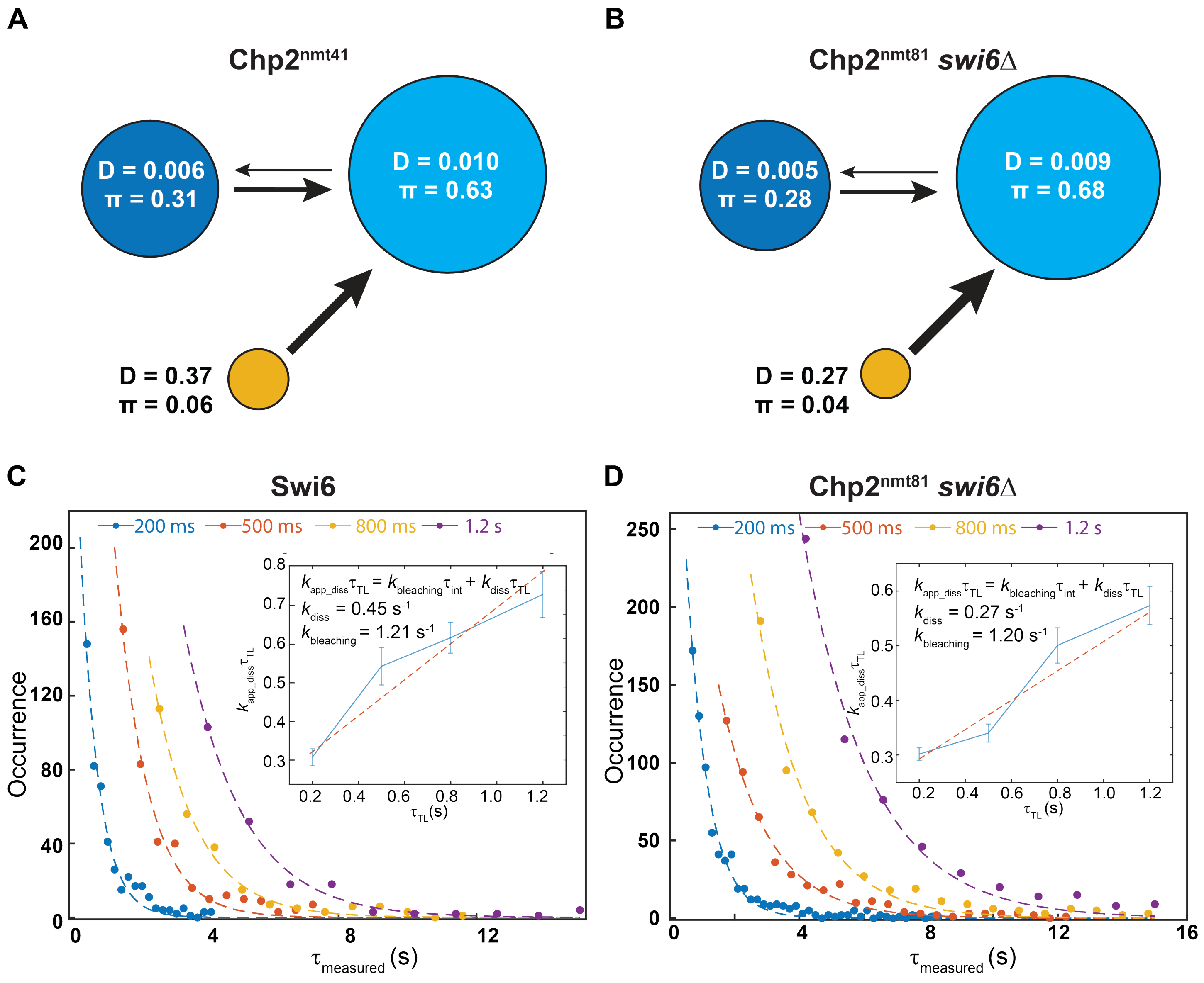


**Supplementary Figure 2. Chp2 has a stronger affinity for H3K9me than Swi6, and the deletion of Swi6 does not affect Chp2-H3K9me binding. A-B**: Inferred transition probabilities between the three mobility states of PAmCherry-Chp2$^{nmt41}$ (**A**) and PAmCherry-Chp2$^{nmt81}$ in *swi6Δ* cells (**B**) from single-molecule tracking with NOBIAS. Diffusion coefficients, *D*, in units of μm$^2$/s and weight fractions, *π*, are indicated. The arrow widths are proportional to the transition probabilities. **C-D**: Dwell time distributions for PAmCherry-Swi6 expressed under its endogenous promoter (**C**) and PAmCherry-Chp2$^{nmt81}$ (**D**) in *swi6Δ* cells. The distributions are shown with fits to an exponential decay. Insert: linear fit (red dashed line) of $k_{app\_diss}\tau_{TL}$ versus $\tau_{TL}$, from which the dissociation rate constant $k_{diss}$ and the photobleaching rate constant $k_{bleaching}$ are obtained. Errors bars are the standard deviation of the exponential decay fitting.

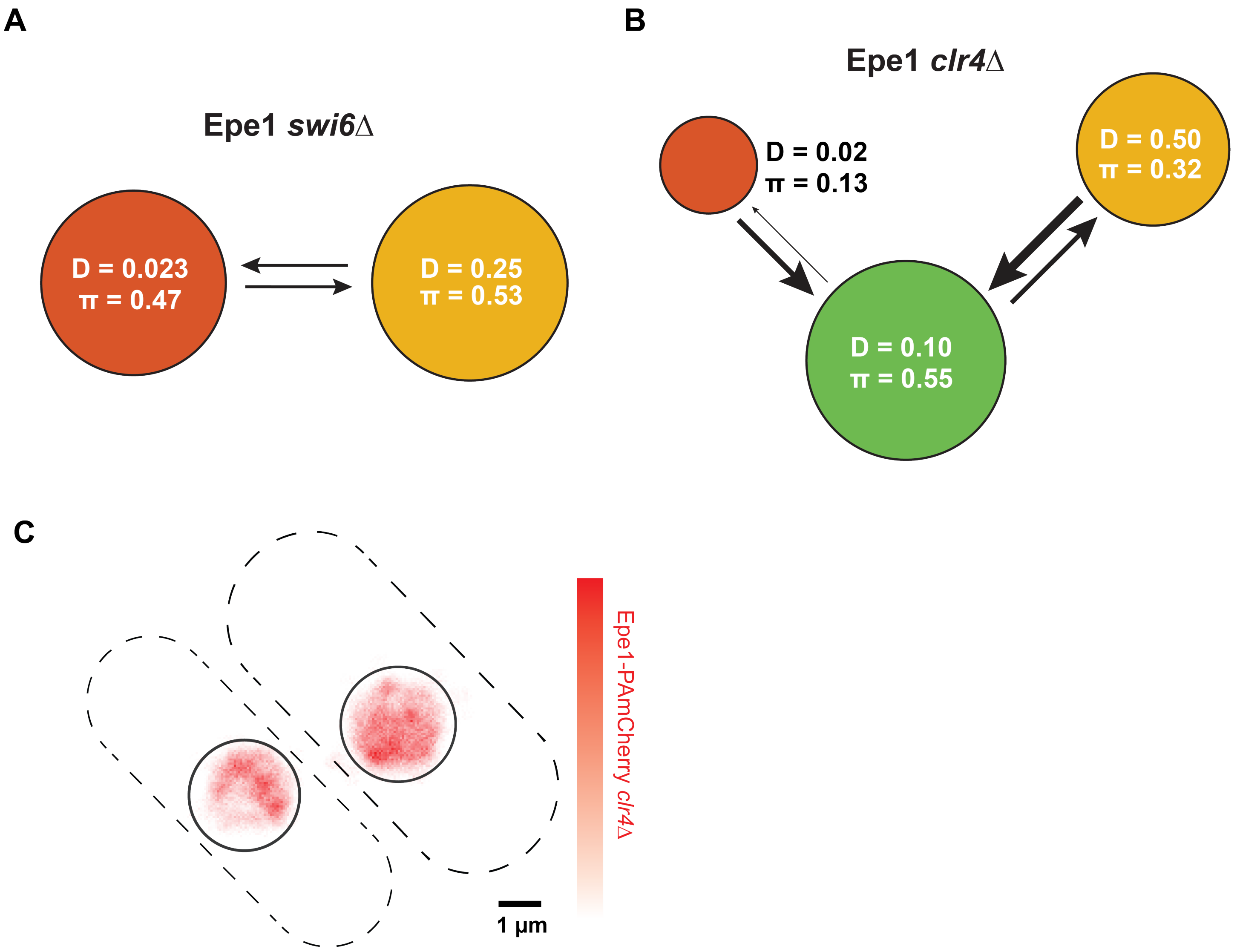


**Supplementary Figure 3. H3K9me reinforces Epe1-Swi6 interactions on-site at heterochromatin and suppresses off-site interactions. A-B**: Inferred probabilities between the mobility states of Epe1-PAmCherry in *swi6Δ* cells (**A**) and in *clr4Δ* cells (**B**) from single-molecule tracking with NOBIAS. The arrow widths are proportional to the transition probabilities. **C**: Reconstructed single-molecule fits density map for Epe1-PAmCherry in *clr4Δ* cells. Dashed lines: approximate *S. pombe* cell outlines; solid circles: approximate nucleus borders.

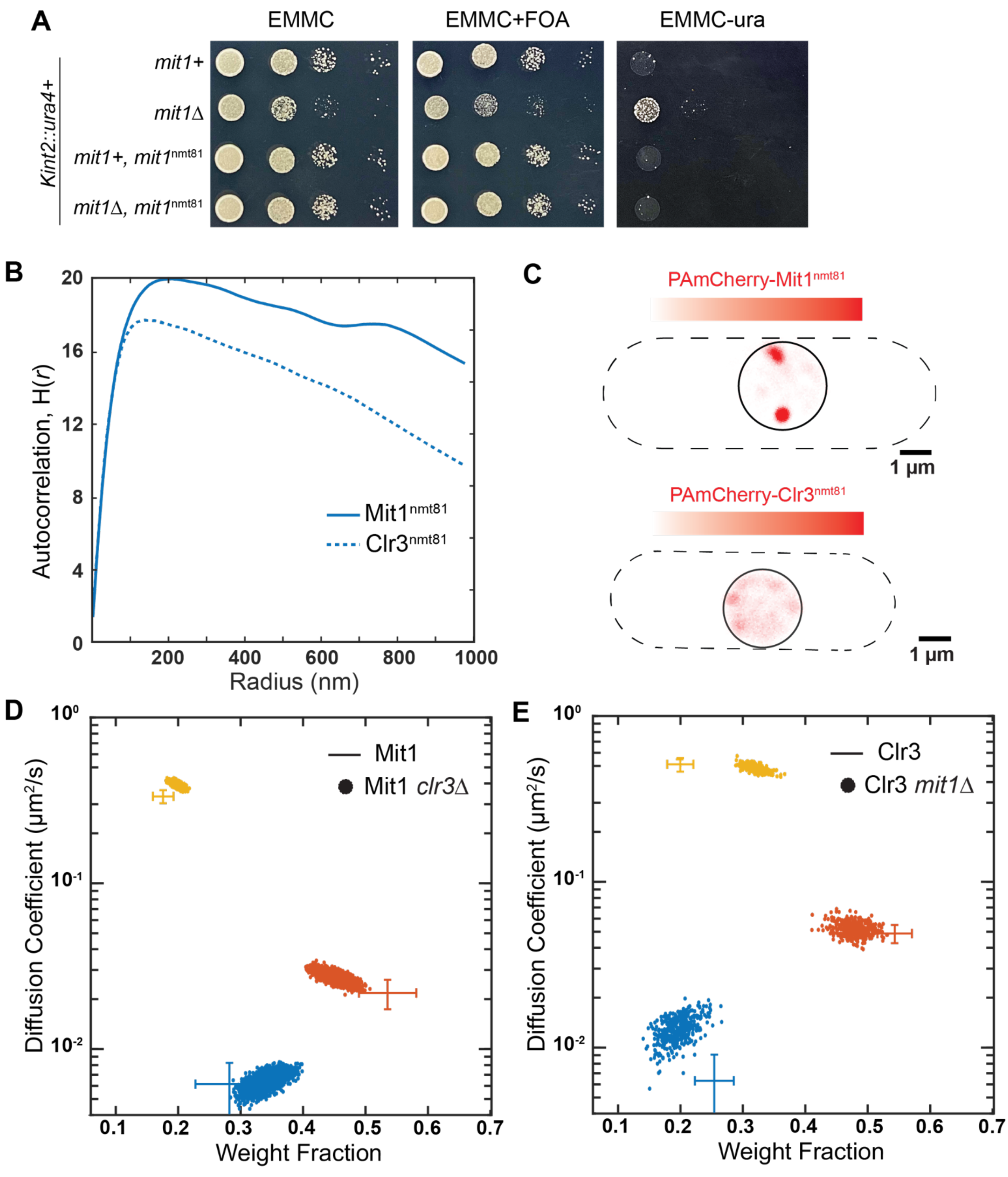


**Supplementary Figure 4. The two SHREC complex subunits proteins Mit1 and Clr3 only assemble at heterochromatin sites (caption on next page).**

**Supplementary Figure 4. The two SHREC complex subunits proteins Mit1 and Clr3 only assemble at heterochromatin sites. A**: Silencing assay using a *ura4*+ reporter inserted at the *mat* locus (*Kint2*:*ura4).* 10-fold serial dilutions of cells were plated on EMMC, EMMC+FOA, and EMM-URA plates to determine if PAmCherry-Mit1$^{nmt81}$ expression can establish heterochromatin in *mit1Δ* cells. **B**: The slow state of Mit1 (solid line) has similar Ripley's H(*r*) value as the slow state of Clr3 (dashed line) in WT cells, indicating that both proteins are clustered in slow states. Each autocorrelation plot is normalized with randomly simulated trajectories from the same state (Methods). **C**: Reconstructed single-molecule fits density map for PAmCherry-Mit1$^{nmt81}$ (top) and PAmCherry-Clr3$^{nmt81}$ (bottom) in WT cells. Dashed lines: approximate *S. pombe* cell outlines; solid circles: approximate nucleus borders. **D:** NOBIAS identifies distinct mobility states for PAmCherry-Mit1$^{nmt81}$ in in *clr3Δ* cells. **E**: NOBIAS also identifies distinct mobility states for PAmCherry-Clr3$^{nmt81}$ in *mit1Δ* cells. Each colored point is the average single-molecule diffusion coefficient of molecules in that state sampled from the posterior distribution of NOBIAS inference at a saved iteration after convergence. The colored crosses show the data from PAmCherry-Mit1$^{nmt81}$ or PAmCherry-Clr3$^{nmt81}$ molecules in WT cells (Figure **5B**).

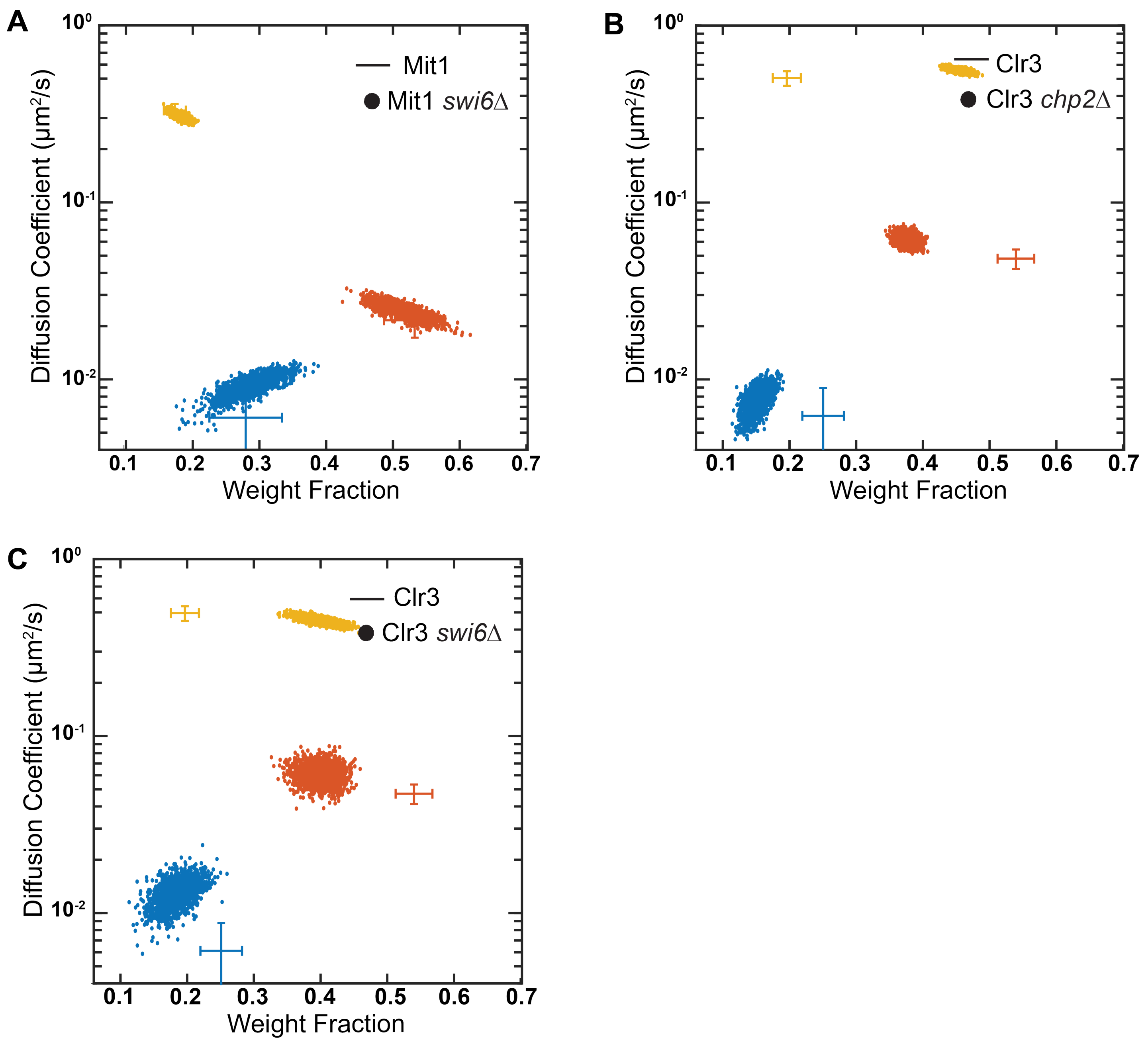


**Supplementary Figure 5. The binding of Clr3 to chromatin partially depends on HP1 and Mit1, while the binding of Mit1 to chromatin does not depend on Clr3. A**: NOBIAS identifies distinct mobility states for PAmCherry-Mit1$^{nmt81}$ in *swi6Δ* cells. **B-C**: NOBIAS also identifies distinct mobility states for PAmCherry-Clr3$^{nmt81}$ in *chp2Δ* cells (**B**) and in *swi6Δ* cells (**C**). Each colored point is the average single-molecule diffusion coefficient of molecules in that state sampled from the posterior distribution of NOBIAS inference at a saved iteration after convergence. The colored crosses show the data from PAmCherry-Mit1$^{nmt81}$ or PAmCherry-Clr3$^{nmt81}$ molecules in WT cells (Figure **5B**).

| *Strain no.* | *Strain genotype* | *Source* | *Related to* |
|---|---|---|---|
| KR2659 | h90 ade6-M210 his2 leu1-32 ura4DS/E kint2::ura4+ chp2Δ::kan | Moazed lab | Figure 1C |
| KR2771 | h90 ade6-M210 his2 leu1-32 ura4DS/E kint2::ura4+ chp2Δ::kan, leu1+:nmt41-PAmCherry-Chp2-ADHt #1 | This study | Figure 1C |
| KR2772 | h90 ade6-M210 his2 leu1-32 ura4DS/E kint2::ura4+ chp2Δ::kan, leu1+:nmt41-PAmCherry-Chp2-ADHt #2 | This study | Figure 1C |
| KR2773 | h90 ade6-M210 his2 leu1-32 ura4DS/E kint2::ura4+ chp2Δ::kan, leu1+:nmt1-PAmCherry-Chp2-ADHt #1 | This study | Figure 1C |
| KR2774 | h90 ade6-M210 his2 leu1-32 ura4DS/E kint2::ura4+ chp2Δ::kan, leu1+:nmt1-PAmCherry-Chp2-ADHt #2 | This study | Figure 1C |
| KR2775 | h90 ade6-M210 his2 leu1-32 ura4DS/E kint2::ura4+ chp2Δ::kan, leu1+:nmt81-PAmCherry-Chp2-ADHt #1 | This study | Figure 1C |
| KR2776 | h90 ade6-M210 his2 leu1-32 ura4DS/E kint2::ura4+ chp2Δ::kan, leu1+:nmt81-PAmCherry-Chp2-ADHt #2 | This study | Figure 1C |
| KR2598 | h90 ade6-M216 leu1+ nmt81-pamcherry-chp2-ura4-D18 | This study | Figure1D |
| KR2616 | h90 leu1+:nmt81-pamcherry-chp2, clr4::KanMX6#1 | This study | Figure 1E |
| KR2662 | h90 ade6-M216 leu1+ nmt1-pamcherry-chp2-ura4-D18 #1 | This study | Figure 1G |
| KR2664 | h90 ade6-M216 leu1+ nmt41-pamcherry-chp2-ura4-D18 #1 | This study | Figure 1F |
| KR2620 | h90 leu1+:nmt81-pamcherry-chp2, swi6Δ::natMX6 #2 | This study | Figure S1C |
| KR1176 | h90 leu1+:mNeonGreen-Swi6 WT-kanMX6 Epe1-PAmcherry hphMX6 | This study | Figure 3AB |
| KR2710 | h90, Epe1-PAmcherryhphMX6, clr4Δ::KanMX6#1 | This study | Figure 3C |
| KR2601 | h90 Epe1-PAmcherryhphMX6, swi6Δ::natMX6 #1 | This study | Figure 3D |
| KR2322 | h90 ade6-M216 leu1+-nmt81-mit1-pamcherry ura4-D18 #10 | This study | Figure 4B |
| KR2597 | h90 ade6-M216 leu1+ nmt81-pamcherry-clr3-ura4-D18 | This study | Figure 4B |
| KR2428 | h90 leu1-32 his2- ura4 DS/E ade6-M210 Kint2::ura4+, leu1+:nmt41-mit1-pamcherry#1 | This study | Figure S4A |
| KR2430 | h90 leu1-32 his2- ura4 DS/E ade6-M210 Kint2::ura4+, mit1Δ, leu1+:nmt41-mit1-pamcherry#1 | This study | Figure S4A |
| KR2432 | h90 leu1-32 his2- ura4 DS/E ade6-M210 Kint2::ura4+, leu1+:nmt81-mit1-pamcherry#1 | This study | Figure S4A |
| KR2434 | h90 leu1-32 his2- ura4 DS/E ade6-M210 Kint2::ura4+, mit1Δ, leu1+:nmt81-mit1-pamcherry#1 | This study | Figure S4A |
| KR2407 | h90 leu1-32 ura4DS/E ade6-M210? otr1R::ura4? mit1-pamcherry-hphMX6 leu1+:nmt81-mit1-pamcherry clr4Δ::KanMX6 #4 | This study | Figure 5A |
| KR2607 | h90 ade6-M216 leu1+ nmt81-pamcherry-clr3-ura4-D18 clr4Δ::KanMX6 #1 | This study | Figure 5B |
| KR2411 | h90 leu1-32 ura4DS/E ade6-M210? otr1R::ura4? mit1-pamcherry-hphMX6 leu1+:nmt81-mit1-pamcherry chp2Δ::kanMX6 #9 | This study | Figure 5E |

| | | | |
|---|---|---|---|
| KR2563 | h90 leu1-32 ura4DS/E ade6-M210? otr1R::ura4? mit1-pamcherry-hphMX6 leu1+:nmt81-mit1-pamcherry chp2Δ::kanMX6 swi6D::natMX6 #1 | This study | Figure 5F |
| KR2404 | h90 leu1-32 ura4DS/E ade6-M210? otr1R::ura4? mit1-pamcherry-hphMX6 leu1+:nmt81-mit1-pamcherry swi6Δ::NATMx6 #1 | This study | FigureS5A |
| KR2416 | h90 leu1-32 ura4DS/E ade6-M210? otr1R::ura4? mit1-pamcherry-hphMX6 leu1+:nmt81-mit1-pamcherry clr1Δ::kanmx6 #2 | This study | FigureS5B |
| KR2604 | h90 ade6-M216 leu1+:nmt81-pamcherry-clr3-ura4-D18 chp2Δ::KanMX #1 | This study | FigureS5C |
| KR2611 | h90 ade6-M216 leu1+:nmt81-pamcherry-clr3-ura4-D18 swi6Δ::natMX6 #2 | This study | FigureS5D |
| KR2412 | h90 leu1-32 ura4DS/E ade6-M210? otr1R::ura4? mit1-pamcherry-hphMX6 leu1+:nmt81-mit1-pamcherry clr3Δ::kanmx6 #1 | This study | FigureS5E |
| KR2615 | h90 ade6-M216 leu1+:nmt81-pamcherry-clr3 ura4-D18 mit1Δ #7 | This study | FigureS5F |
